# Constitutive Yap activation in distal nephron segments disrupts epithelial identity and nephron patterning

**DOI:** 10.64898/2025.12.10.693519

**Authors:** Zeinab Dehghani-Ghobadi, Eunah Chung, Mohammed Sayed, Christopher Ahn, Hyojin Alex Choi, Annissa Aamoum, Benjamin R. Thomson, Yueh-Chiang Hu, Hee-Woong Lim, Joo-Seop Park

## Abstract

The distal nephron segments play a critical role in maintaining electrolyte balance, yet the mechanisms that preserve epithelial identity and segmental organization within this region remain poorly defined. Yes-associated protein (Yap), a key effector of Hippo signaling, is essential for kidney development, but its function in distal nephron epithelia is unknown. Using a genetic gain-of-function approach to activate Yap selectively in distal nephron segments, we found that sustained Yap activity profoundly disrupts epithelial organization and nephron patterning. Lineage tracing revealed that both distal convoluted tubule and connecting tubule cells originate from *Slc12a3*-expressing cells, and Yap activation in these segments led to increased proliferation, displacement of lineage-labeled cells beyond expected segment boundaries, and loss of segment-specific gene expression. These changes were accompanied by defects in apicobasal polarity and junctional integrity, consistent with epithelial plasticity. Unexpectedly, Yap activation in distal nephron segments also suppressed proximal tubule gene expression, indicating non-cell-autonomous effects on nephron differentiation. Together, these findings identify Yap as a critical regulator of epithelial identity in the distal nephron segments and reveal a previously unrecognized role for Hippo signaling in coordinating intersegmental organization during kidney development.

## INTRODUCTION

The kidney maintains systemic homeostasis by regulating electrolyte balance, fluid volume, and blood pressure (1). The nephron, the functional unit responsible for these tasks, consists of two main parts: the renal corpuscle, which contains podocytes and parietal epithelial cells, and the tubular system, which includes the proximal convoluted tubule, loop of Henle, distal convoluted tubule (DCT), and connecting tubule (CNT) (1, 2). Dysfunction in any of these segments can impair kidney function, resulting in electrolyte disorders, interstitial fibrosis, and chronic kidney disease (3).

Among tubular segments, the DCT plays a critical role in the fine-tuning of electrolyte reabsorption. Positioned between the thick ascending limb of the loop of Henle and the CNT, the DCT consists of two functionally distinct segments: DCT1 and DCT2 (4, 5). Both segments contribute to sodium and chloride reabsorption through the sodium-chloride cotransporter (Slc12a3, also known as NCC). While DCT1 mediates magnesium homeostasis, DCT2 regulates calcium and potassium transport and exhibits increased sensitivity to aldosterone (4, 5).

The Hippo signaling pathway is an evolutionarily conserved key regulator of cell proliferation, differentiation, and organ size (6–9). Its downstream effectors are the transcriptional coactivators Yap (Yes-associated protein 1; *Yap1*) and Taz (WW domain-containing transcription regulator 1; *Wwtr1*). In mammals, activation of the Hippo pathway triggers a kinase cascade involving MST1/2 and LATS1/2, which phosphorylate Yap and Taz. This phosphorylation leads to their cytoplasmic retention and degradation. When the Hippo pathway is inactive, unphosphorylated Yap and Taz translocate into the nucleus, where they interact with TEAD transcription factors to drive the expression of their target genes.

During kidney development, Hippo⍰Yap signaling plays essential roles during early nephron morphogenesis, with nuclear Yap detected in epithelial compartments of the S⍰shaped body, including the distal domain (10). As the distal S⍰shaped body represents the epithelial progenitor population for the distal convoluted tubule and connecting tubule (11), these observations are consistent with a potential role for Yap in early distal lineage formation. However, how Hippo⍰Yap signaling is regulated as distal epithelial cells transition from progenitor states to terminal differentiation remains unclear. Given the extensive epithelial remodeling that accompanies late nephrogenesis and early postnatal maturation, it is plausible that Yap activity requires precise temporal regulation during distal tubule differentiation, yet whether repression of Yap signaling is required to preserve distal epithelial identity and structural maturation remains unknown.

In contrast to these early epithelial contexts, Hippo⍰Yap signaling has been examined primarily in mesenchymal populations, including nephron progenitors and stromal lineages (12, 13). Moreover, studies investigating Yap function in renal epithelial cells have largely focused on adult pathological conditions such as injury, fibrosis, cystic disease, or oncogenesis (14–19), leaving its role in differentiated epithelial nephron segments during kidney development, after segmental identity has been established, relatively unexplored.

In this study, we used the *Slc12a3-IRES-Cre* driver to activate a constitutively active Yap allele (*Col1a1-Yap5SA*) specifically in distal nephron segments. Through epithelial lineage tracing and analysis of segment-specific markers, we examined how sustained Yap activity influences tubular identity, cell polarity, and nephron organization. By activating Yap after distal tubule specification, this model allows interrogation of Hippo pathway repression in post⍰specification epithelial maintenance, while avoiding perturbation of early nephron induction or proliferative nephrogenesis. This model provides a platform to dissect both cell-intrinsic and non-cell-autonomous consequences of Hippo pathway inhibition within the distal nephron.

## RESULTS

### DCT cells give rise to CNT cells during nephron development

The DCT of the nephron can be targeted using the previously reported *Slc12a3-IRES-CreERT2* allele (20). However, because nephrogenesis occurs in waves during development, DCT cells are generated at different times, making it impractical for a tamoxifen⍰inducible system to achieve complete and uniform targeting of all DCT cells. To overcome this limitation, we generated *Slc12a3-IRES-Cre* (hereafter *Slc12a3Cre*) using CRISPR/Cas9 genome editing to convert *Slc12a3-IRES-CreERT2* into a constitutively active allele. We assessed the recombinase activity of this new Cre line by lineage tracing with a *Rosa26-Sun1* reporter (21). As expected, all Slc12a3+ cells were positive for GFP, confirming that *Slc12a3Cre* efficiently targets the entire DCT segment, including DCT1, the transition zone, and DCT2 (Figure 1A; Supplemental Figures 1A and 2). Unexpectedly, we also observed GFP+ cells that lacked Slc12a3. Immunofluorescence analysis with distal nephron markers showed that these GFP+ Slc12a3-cells expressed S100g, suggesting that they are the CNT segment (Figure 1A; Supplemental Figures 1A and 2). This observation suggests that CNT cells arise from the DCT lineage during development.

**Figure 1.**
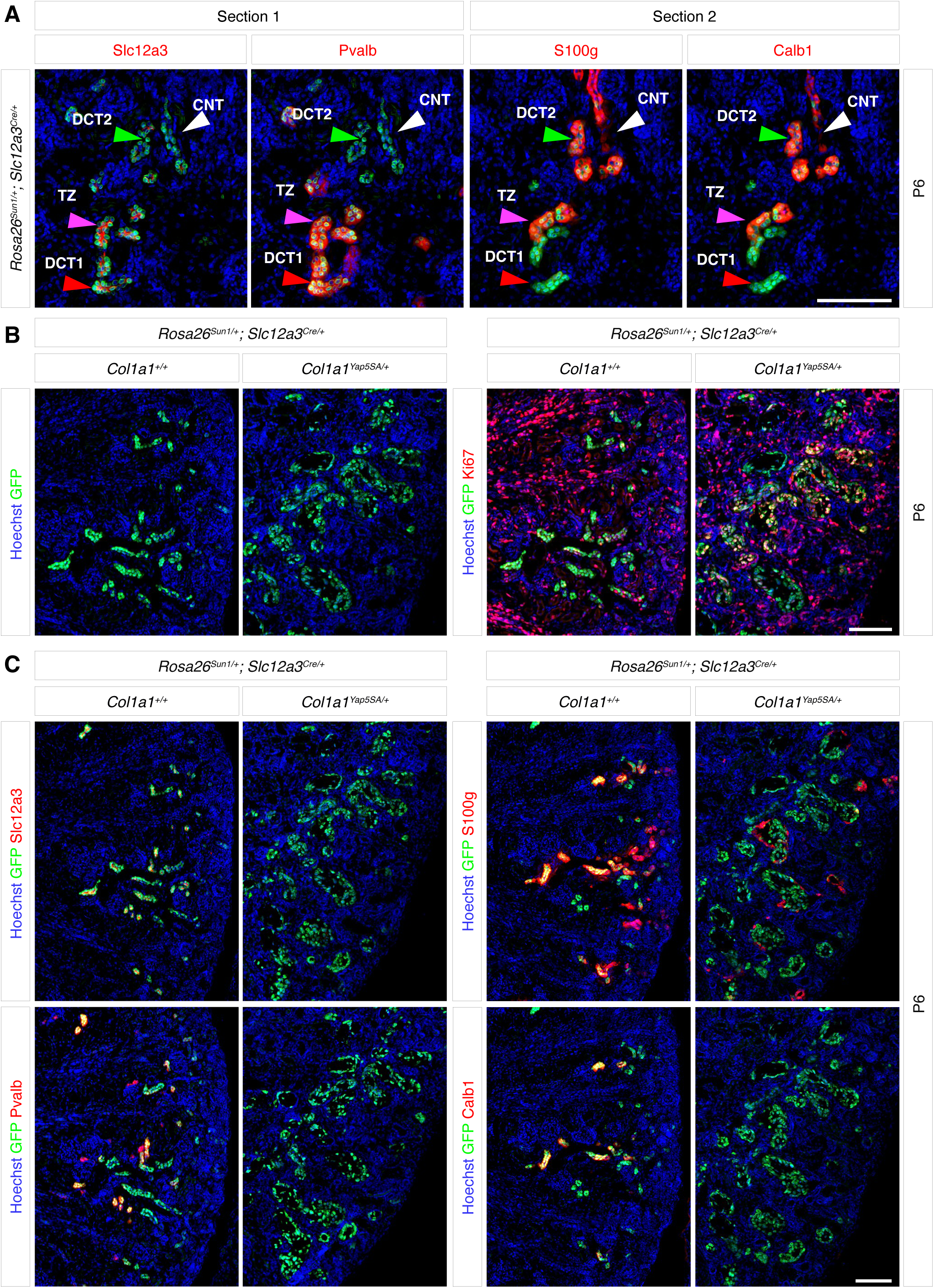
Constitutive activation of Yap in the distal convoluted tubule (DCT) and connecting tubule (CNT) leads to increased proliferation and disrupts segmental identity (A) Lineage tracing using the *Rosa26-Sun1* reporter (nuclear membrane GFP) shows that *Slc12a3Cre* targets both DCT and CNT. Slc12a3 marks both DCT1 and DCT2, while Pvalb marks DCT1 only. S100g and Calb1 mark DCT2 and CNT. The transition zone (TZ) is marked by both DCT1 and DCT2 markers (see Supplemental Figure 2). Adjacent sections of the same kidney are shown. (B) In control kidneys, only a small subset of DCT and CNT cells (GFP+) are positive for Ki67, indicating low proliferative activity. In contrast, DCT and CNT cells with persistent Yap activation show strong Ki67 staining, consistent with increased proliferation. (C) GFP+ cells in the control kidney express markers characteristic of distal nephron segments, while GFP+ cells in the mutant kidney lack these markers, suggesting disrupted segmental identity. (A-C) Representative images from three independent experiments are shown. Stage, postnatal day 6; Scale bar, 100 µm

**Figure 2.**
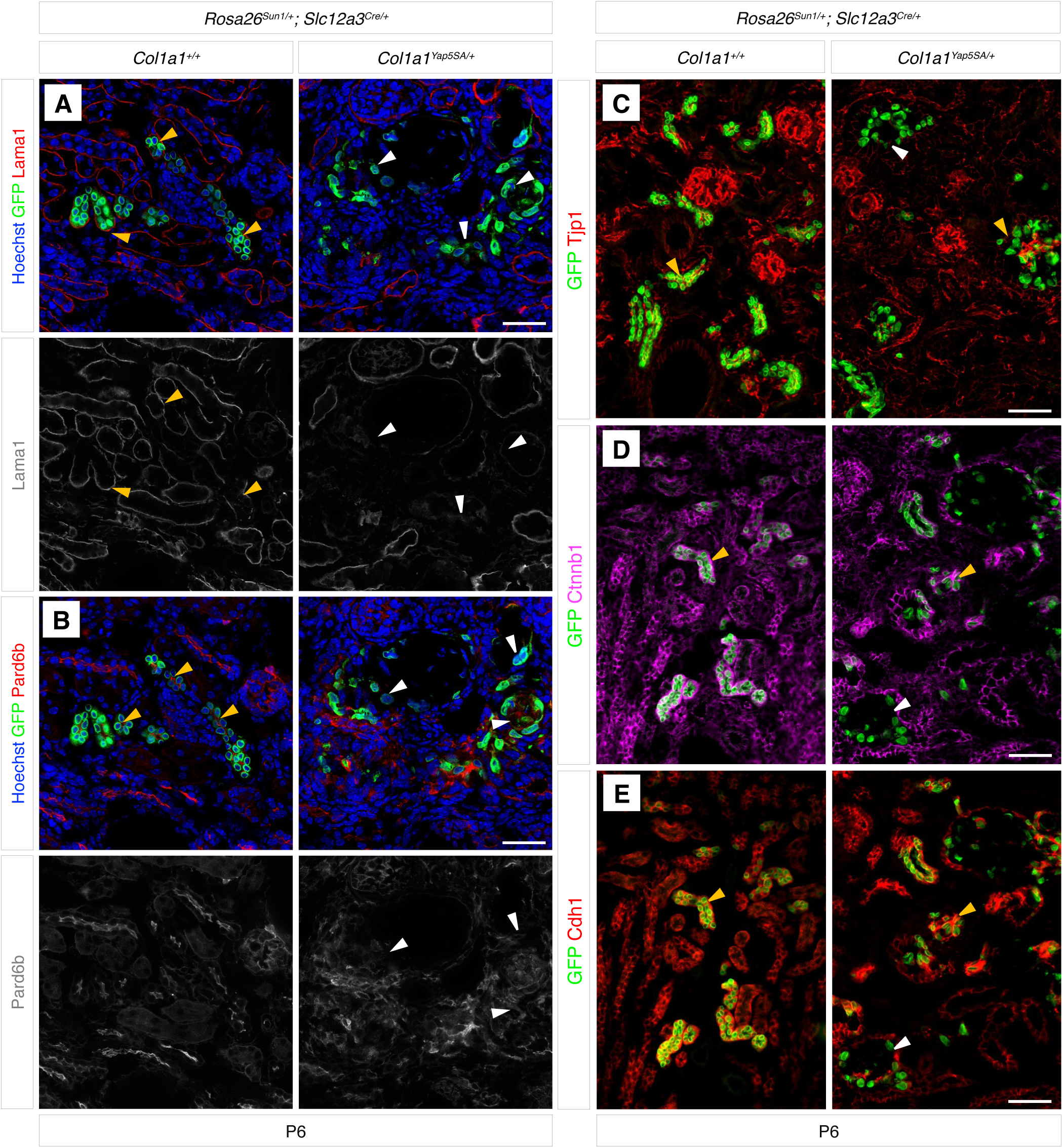
Constitutive activation of Yap in DCT and CNT impairs epithelial polarity and junctional organization. (A, B) In the control kidney, the basal and apical sides of GFP+ cells are marked by Lama1 and Pard6b, respectively (marked by orange arrowheads). In the mutant kidney, GFP+ cells (marked by white arrowheads) lack basal Lama1 and apical Pard6b. (C-E) In the control kidney, GFP+ cells express the tight junction marker *Tjp1* (*ZO1*) and the adherens junction markers, such as Ctnnb1 and Cdh1 (marked by orange arrowheads). In mutant kidneys, GFP+ cells exhibit either a loss (white arrowheads) or mislocalization (orange arrowheads) of junctional markers, suggesting impaired epithelial integrity. (A-E) Representative images from three independent experiments are shown. Stage, postnatal day 6; Scale bar, 100 µm

### Constitutive Yap activation in DCT and CNT cells increases proliferation and disrupts segmental identity

To establish the physiological context for Yap regulation in the distal nephron, we first examined Yap subcellular localization during early postnatal kidney development. Immunofluorescence analysis revealed that at postnatal day 1, both proximal tubule and DCT/CNT cells exhibit nuclear Yap, whereas by postnatal day 6 nuclear Yap is retained in proximal tubule cells but is largely absent from DCT/CNT cells (Supplemental Figure 3). These observations indicate that Yap activity is normally downregulated in distal tubule cells during postnatal maturation, defining a developmental stage at which sustained Yap activation would be non-native to distal nephron epithelia.

To test the consequences of preventing this developmental repression of Yap activity, we generated a Cre-dependent Yap gain-of-function (GOF) allele (Supplemental Figure 4). Upon Cre-mediated inversion, this allele expresses *Yap5SA*, a modified form in which five serine residues are replaced with alanine (22, 23). These substitutions prevent phosphorylation by Hippo kinases, thereby blocking Yap inactivation and rendering it constitutively active. Using CRISPR/Cas9 genome editing, we inserted the Cre-inducible *Yap5SA* cassette into the safe-harbor *Col1a1* locus (hereafter *Col1a1-Yap5SA*; Supplemental Figure 4). Because constitutive Yap activation mimics Hippo pathway inhibition regardless of upstream inputs, this allele provides a powerful tool to assess the effects of sustained Yap activity in vivo.

To investigate the role of Hippo signaling in the distal nephron segments, we crossed *Slc12a3Cre* with *Col1a1-Yap5SA* to generate Yap GOF mutant kidneys, in which Yap is constitutively active in the DCT and CNT. A *Rosa26-Sun1* reporter was included to mark cells that had experienced Cre-mediated recombination. Because the Yap GOF allele encodes a 3xFLAG⍰tagged Yap protein, we first confirmed transgene expression within the targeted cell population by FLAG immunofluorescence. In mutant kidneys, all GFP+ distal nephron epithelial cells were FLAG⍰positive, confirming robust Yap transgene expression in Slc12a3⍰lineage cells (Supplemental Figure 5).

At postnatal day 6 (P6), mutant kidneys contained markedly increased numbers of GFP+ cells compared to controls, often arranged in multiple layers surrounding dilated lumens (Figure 1B, left panel; Supplemental Figure 1B). These GFP+ cells exhibited a significant increase in Ki67 staining, indicating hyperproliferation associated with constitutive Yap activation (Figure 1B, right panel; Supplemental Figure 1B). Immunofluorescence analysis using distal nephron markers revealed distinct segment⍰specific expression patterns in control kidneys but a near⍰complete loss of expression in mutant kidneys, consistent with disruption of distal nephron identity (Figure 1C; Supplemental Figure 1C, D). S100g expression was still detected in mutant kidneys; however, the majority of S100g+ cells were GFP⍰negative, suggesting that they had not undergone sustained Yap activation (Figure 1C; Supplemental Figure 1D).

To determine the developmental onset of the phenotype, we next examined kidneys at embryonic day 18.5 (E18.5) and postnatal day 3 (P3). At E18.5, mutant kidneys exhibited early and heterogeneous defects in distal nephron identity, with subsets of GFP+ tubules lacking expression of distal segment markers (Supplemental Figure□6). By P3, loss of DCT and CNT marker expression became widespread among GFP+ cells, indicating a robust and uniform disruption of distal nephron segment identity (Supplemental Figure□7).

Hematoxylin and eosin staining further demonstrated pronounced tubular dilation and under⍰developed papillary architecture in mutant kidneys (Supplemental Figure 8A). Kaplan-Meier survival analysis revealed markedly increased neonatal mortality, with no mutant animals surviving beyond postnatal day 7 (Supplemental Figure 8B).

Together, these data demonstrate that sustained Yap activation in distal nephron epithelial cells leads to early disruption of epithelial identity, followed by hyperproliferation, architectural disorganization, and early postnatal lethality.

### Constitutive Yap activation disrupts polarity and junctional organization in DCT and CNT cells

Our observation that GFP+ cells in the mutant kidney often failed to remain as a monolayer prompted us to examine how constitutive Yap activation affects epithelial polarity and junctional organization. In control kidneys, GFP+ tubules showed Lama1 at the basal side and Pard6b at the apical side (Figure 2A-B). In contrast, GFP+ cells in mutant kidneys showed a marked loss of basal Lama1 and disrupted apical distribution of Pard6b, indicating impaired polarity (Figures 2A-B;).

To determine when these polarity defects first arise during development, we examined distal nephron tubules at E18.5 and P3. At both stages, mutant kidneys exhibited heterogeneous polarity defects among GFP+ tubules, with some retaining polarized Lama1 and Pard6b localization and others showing clear loss of polarity (Supplemental Figure 9A). By P3, a greater proportion of GFP+ tubules displayed polarity defects compared with E18.5, indicating a progressive loss of epithelial polarity during postnatal maturation (Supplemental Figure 9B).

Furthermore, in GFP+ cells in mutant kidneys, junctional proteins such as Tjp1 (ZO1), Ctnnb1 (b-catenin), and Cdh1 (E-cadherin) were frequently absent or mislocalized from their normal junctional positions (Figures 2C-E; Supplemental Figure 10), suggesting destabilization of cell-cell adhesion complexes. These alterations are consistent with Yap’s established role in maintaining epithelial architecture in other tissues (9, 24) and likely contribute to the loss of segment-specific identity in Yap GOF mutant kidneys.

### Constitutive Yap activation impairs distal nephron identity and alters epithelial spatial organization

Gata3, a key transcription factor regulating the branching of the collecting duct (CD) during development (25, 26), is expressed in DCT, CNT, and CD of the renal epithelia (2). In Yap GOF mutant kidneys, Gata3 was largely absent in DCT and CNT cells marked by activation of the *Rosa26-Sun1* reporter (Figure 3A; Supplemental Figure 11), indicating that sustained Yap activity disrupts segmental identity and differentiation in the distal nephron.

**Figure 3.**
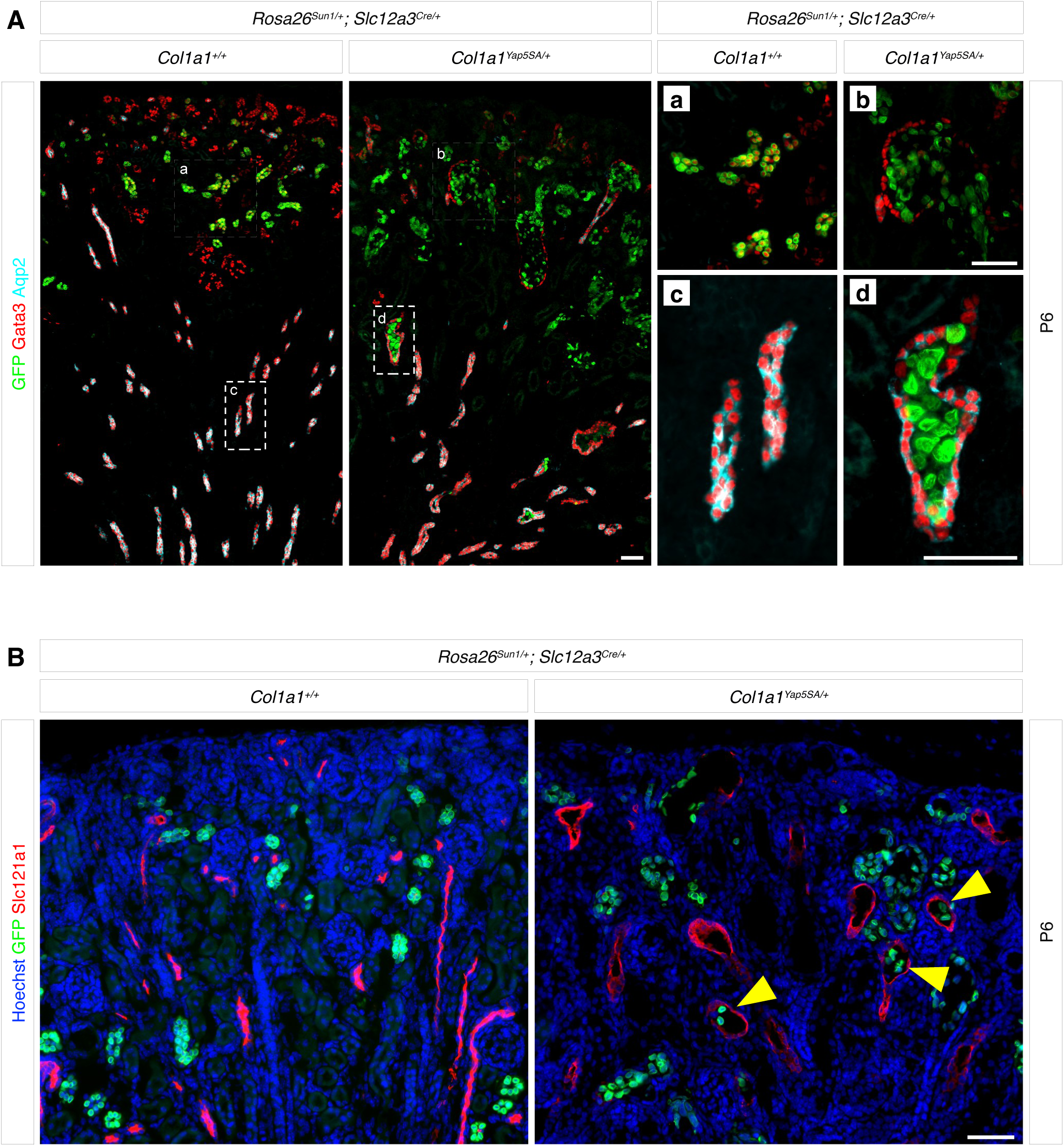
Constitutive Yap activation in DCT and CNT is associated with displacement of lineagelZI labeled cells beyond normal segmental boundaries In both control and mutant kidneys, *Slc12a3Cre* activates GFP expression in DCT and CNT, but not in the collecting duct (CD). (A) The left panel shows a low-magnification view of the kidney, with the regions outlined by dotted squares shown at higher magnification in the right panel (a-d). *Aqp2* marks CD while Gata3 marks DCT, CNT, CD, and mesangial cells. (a,b) GFP+ cells in the control kidney are positive for *Gata3*, while those in the mutant kidney lack Gata3. (c,d) In the mutant kidney, GFP+ cells are located within the lumen of Aqp2+ Gata3+ CD cells. (B) Slc12a1 marks the ascending limb of the loop of Henle. In the mutant kidney, GFP+ cells are located within the lumen of Slc12a1+ cells (marked by yellow arrowheads). Representative images from three independent experiments are shown. Stage, postnatal day 6; Scale bar, 50 µm.

In addition to loss of segmental identity, GFP⍰positive lineage⍰labeled distal epithelial cells were detected outside their expected segmental locations. Rosa26⍰Sun1 reporter⍰positive Yap GOF cells were observed within collecting duct structures marked by Gata3 and Aqp2 (Figure 3A; Supplemental Figure 11). GFP+ reporter cells were also identified within the thick ascending limb of the loop of Henle, as marked by Slc12a1 (Figure 3B), indicating altered spatial localization of distal nephron-derived cells relative to normal segmental boundaries.

The macula densa (MD) plays a central role in stimulating renin (*Ren1*) release from the juxtaglomerular apparatus (JGA) by sensing tubular NaCl delivery and producing paracrine mediators such as prostaglandin E_2_ (via COX-2) and nitric oxide (27). Because the MD represents a specialized subset of the thick ascending limb of the loop of Henle, it can be identified as Slc12a1+ cells positioned adjacent to the glomerulus (Supplemental Figure 12A). In Yap GOF mutant kidneys, GFP+ cells were observed within the lumen of the MD, indicating aberrant localization of distal nephron-derived cells within this specialized segment (Supplemental Figure 12A).

Consistent with disruption of MD-JGA signaling, *Ren1* expression was absent in the mutant kidneys (Supplemental Figure 12B). In parallel, RNA⍰seq analysis revealed coordinated downregulation of multiple components of the renin⍰angiotensin system, including *Ren1*, *Ace*, *Agtr1a*, and *Agtr2*, in Yap GOF kidneys (Supplemental Table□1). Together, these findings indicate that constitutive Yap activation disrupts distal nephron organization and is associated with impaired MD⍰dependent endocrine signaling, providing a potential mechanistic link between distal epithelial disorganization and suppression of the MD⍰JGA axis.

To determine whether disruption of the MD⍰JGA axis is associated with impaired nephron flow, we performed an in vivo fluorescent dextran delivery assay at P3 (28). The MD⍰JGA axis is required for maintaining normal glomerular filtration by regulating renin release, intrarenal perfusion, and downstream tubular delivery. Following intravenous administration of 10⍰kDa Texas Red⍰conjugated dextran, control kidneys showed robust luminal dextran signal throughout nephron tubules, consistent with effective filtration and downstream delivery (Supplemental Figure□13). In contrast, mutant kidneys exhibited markedly reduced dextran signal, approaching background levels observed in non⍰injected controls (Supplemental Figure 13). These findings indicate that luminal delivery of filtrate is severely impaired in Yap GOF kidneys, consistent with a functional disruption of nephron flow along the tubular axis.

### Kidney-wide transcriptomic responses to constitutive Yap activation in distal nephron segments

To evaluate the impact of constitutive Yap activation in DCT and CNT on kidney-wide transcriptional programs, we performed bulk RNA-seq analysis on P3 kidneys from mutants and littermate controls (Supplemental Table 1). Our analysis identified 2,310 differentially expressed genes (DEGs), including 1,116 upregulated and 1,194 downregulated transcripts (FC > 1.5, FDR < 0.05) (Supplemental Figure 14A). To determine the cellular origins of these DEGs, we mapped them onto our previously published single-cell RNA-seq dataset of developing mouse kidneys (Supplemental Figure 14B). In the UMAP projection, downregulated genes were enriched in clusters corresponding to the proximal tubule, thick ascending limb, and DCT (Supplemental Figure 14C). In contrast, upregulated genes were predominantly associated with clusters representing immune cell populations (Supplemental Figure 14D). Consistent with these assignments, gene ontology analysis showed that downregulated genes were strongly enriched for proximal tubule functions, including amino-acid catabolism, fatty-acid β-oxidation, and brush-border organization (Supplemental Figure 15A). In contrast, upregulated genes were enriched for inflammatory and immune-activation pathways, including TNF, MAPK, and NF-κB signaling, suggesting that Yap activation induces pro-inflammatory and stress-responsive transcriptional programs. Genes related to cell adhesion, cytoskeletal remodeling, and membrane-raft assembly were also elevated, consistent with altered epithelial architecture and cell–cell interactions in Yap GOF mutant kidneys (Supplemental Figure 15B).

Figure 4 highlights three groups of DEGs and their cell-type-specific expression across the kidney: (1) downregulated DCT/CNT genes, (2) downregulated proximal tubule genes, and (3) upregulated injury-associated genes. Consistent with our findings of the loss of multiple DCT/CNT markers in the Yap GOF mutant kidneys (Figure 1C; Supplemental Figure 1C), the bulk RNA-seq confirmed their downregulation in the Yap GOF mutant kidneys (Figure 4).

**Figure 4.**
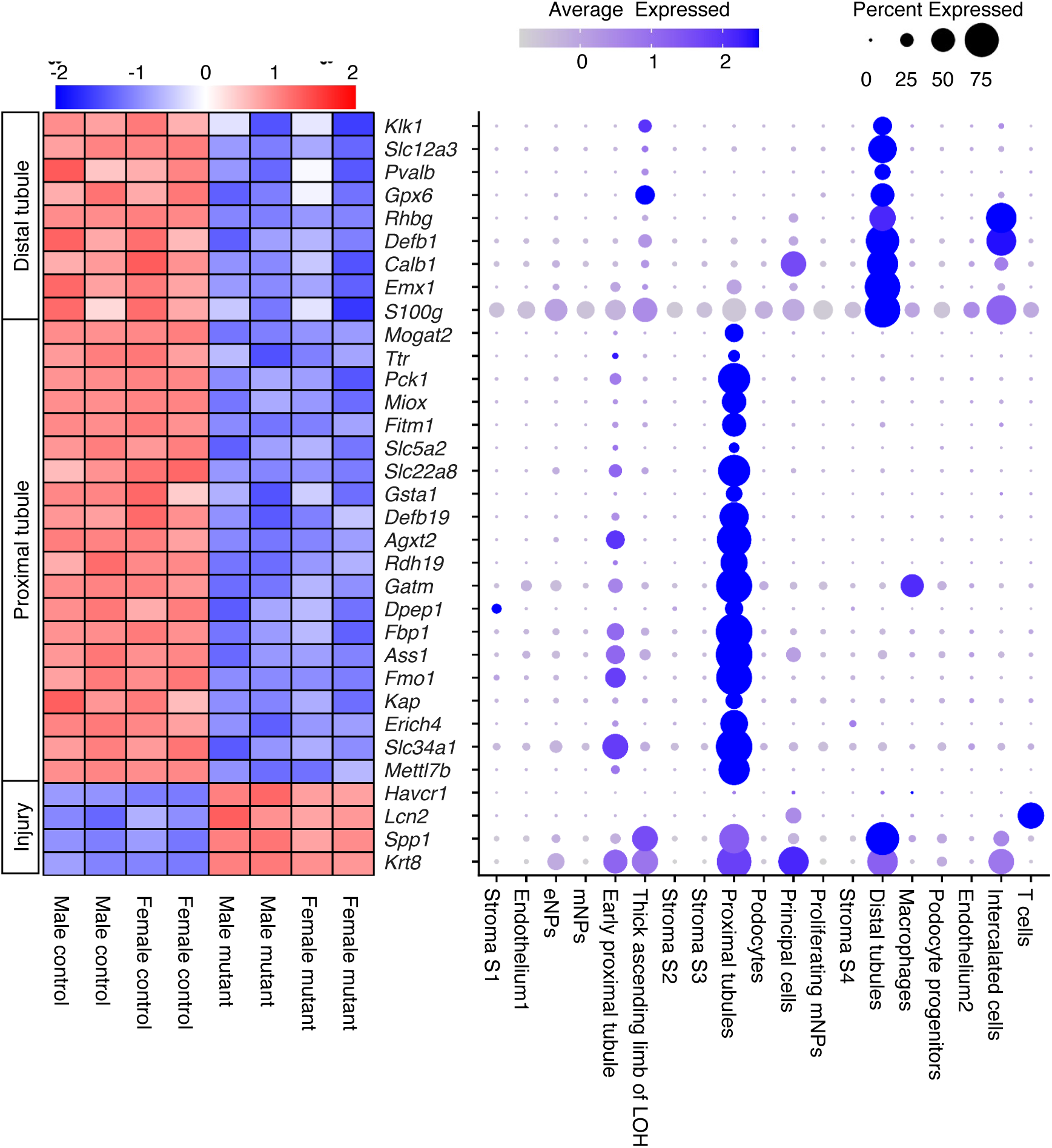
Differential gene expression analysis. Bulk RNA-seq of whole kidneys at postnatal day 3 reveals differentially expressed genes (DEGs) between mutant and control kidneys, as shown in the heatmap. In the mutant kidney, genes normally expressed in distal tubules and proximal tubules are downregulated, while injury-associated genes are upregulated. These expression changes occur regardless of sex. Color intensity represents the magnitude of expression change, with blue indicating decreased and red indicating increased expression. The corresponding dot plot displays cell type-specific expression of marker genes based on previously published scRNA-seq data from mouse kidneys at E18.5 or P0 (GSE214024 and GSE275601). Dot color indicates the average expression level of each gene within a given cell type, while dot size represents the percentage of cells expressing that gene.

We next validated the downregulation of proximal tubule genes in the Yap GOF mutant kidneys by immunofluorescence. Control kidneys showed strong *Lotus tetragonolobus* lectin (LTL) staining, reflecting the abundant glycoproteins in the brush border of mature proximal tubules (Figure 5). In contrast, mutant kidneys exhibited markedly reduced LTL labeling, indicating the absence of mature proximal tubules. Notably, Hnf4a+ proximal tubule cells were still present in mutant kidneys, suggesting that proximal tubule specification still occurred. However, these cells failed to develop into mature proximal tubules. Consistent with impaired maturation, mature proximal tubule markers such as *Pck1*, *Fbp1*, and *Slc5a2* were nearly absent in mutant kidneys, and other proximal tubule markers were also reduced, though to a lesser degree.

**Figure 5.**
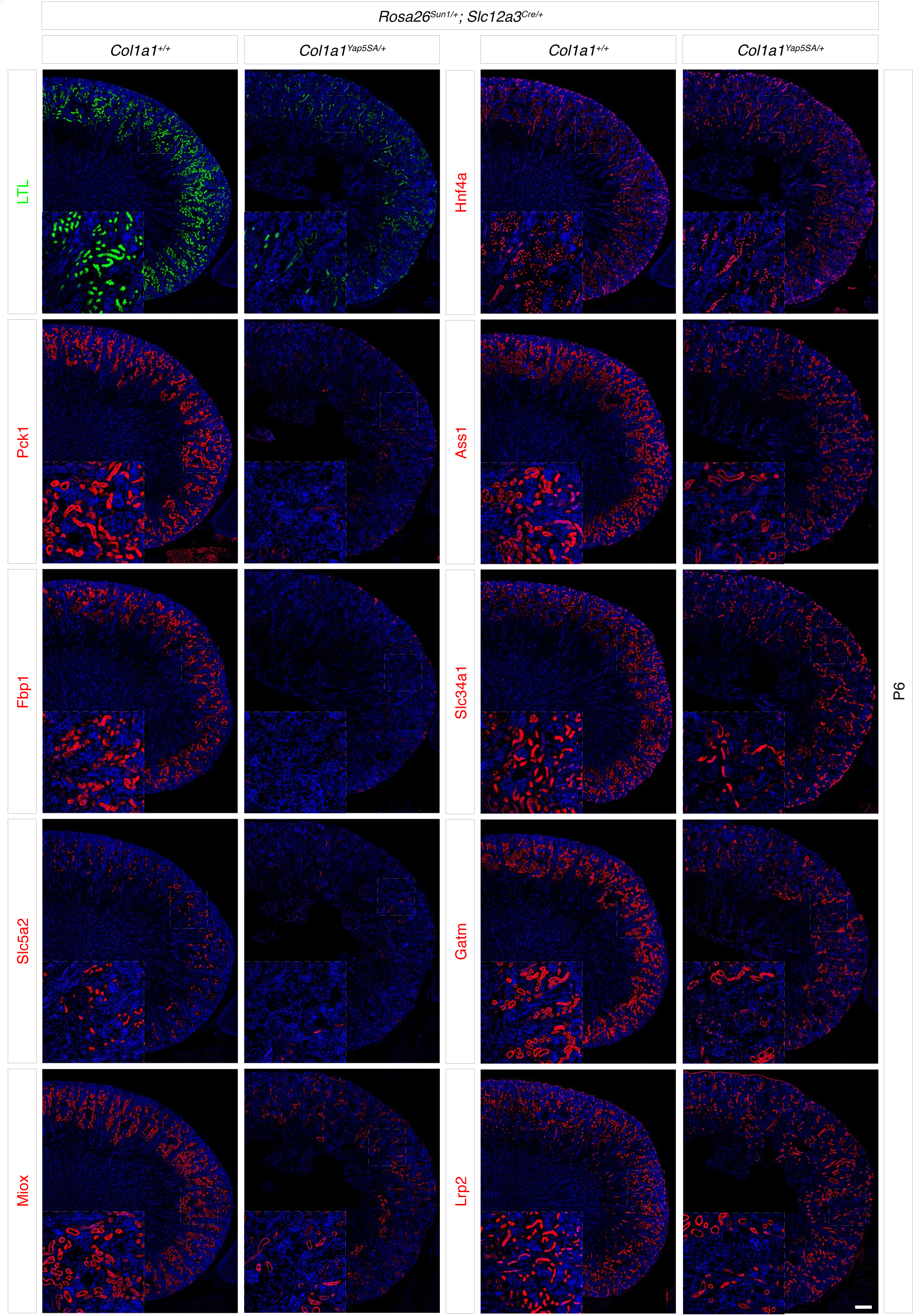
Constitutive Yap activation in DCT and CNT leads to the downregulation of proximal tubule genes. In the mutant kidney, LTL (lotus tetragonolobus lectin) staining is greatly reduced compared with controls, indicating an absence of mature proximal tubules. Expression of proximal tubule markers, such as *Pck1*, *Slc5a2*, and *Fbp1*, is almost absent in the mutant, along with the downregulation of additional proximal tubule genes. Inset panels, denoted by the white dotted squares, present higher⍰magnification views of the specified cortical regions outlined in the overview images. Representative images from three independent experiments are shown. Stage, P6; Scale bar, 200 µm

To determine the developmental onset of this proximal tubule maturation defect, we examined kidneys at E18.5 and P3 (Supplemental Figure 16). At E18.5, mutant kidneys showed preserved expression of proximal tubule markers (Hnf4a, Fbp1, and Pck1), accompanied by weaker LTL staining. By P3, mutant kidneys exhibited broader loss of proximal tubule marker expression and markedly diminished LTL signal, despite the continued presence of Hnf4a⍰positive cells. These findings indicate that proximal tubule specification is initiated but maturation progressively fails during late embryonic and early postnatal development in Yap GOF kidneys.

In addition to proximal tubule defects, the loop of Henle and collecting duct also exhibited structural abnormalities. Segment-specific marker analysis demonstrated impaired tubular segmentation, with markedly reduced expression of *Aqp1* (descending limb), *Slc12a1* (thick ascending limb), and *Aqp2* (collecting duct principal cells) in mutant kidneys (Supplemental Figure 17). Taken together, these results suggest that Yap overactivation in the distal nephron segments blocks proximal tubule maturation, leading to a broad disruption of nephron patterning.

We found that Injury-associated genes, including *Havcr1* (Kim1), *Spp1*, and *Lcn2* (NGAL), were markedly upregulated in mutant kidneys (Figure 4). These transcriptomic changes were corroborated by immunofluorescence staining, which showed increased *Havcr1* and *Spp1* expression in proximal tubules, along with elevated *Krt19* expression in distal nephron segments (Figure 6). Consistent with our observation that immune cell clusters in scRNA-seq data show the enrichment of the genes upregulated in the Yap GOF mutant kidneys (Supplemental Figure 14D), the mutant kidneys exhibited enhanced macrophage infiltration around GFP+ Yap GOF mutant cells (Figure 6), suggesting that immune cell recruitment accompanies the injury response triggered by constitutive Yap activation. Together, these findings indicate that aberrant Yap signaling compromises nephron patterning and promotes inflammatory and cytoskeletal remodeling responses.

**Figure 6.**
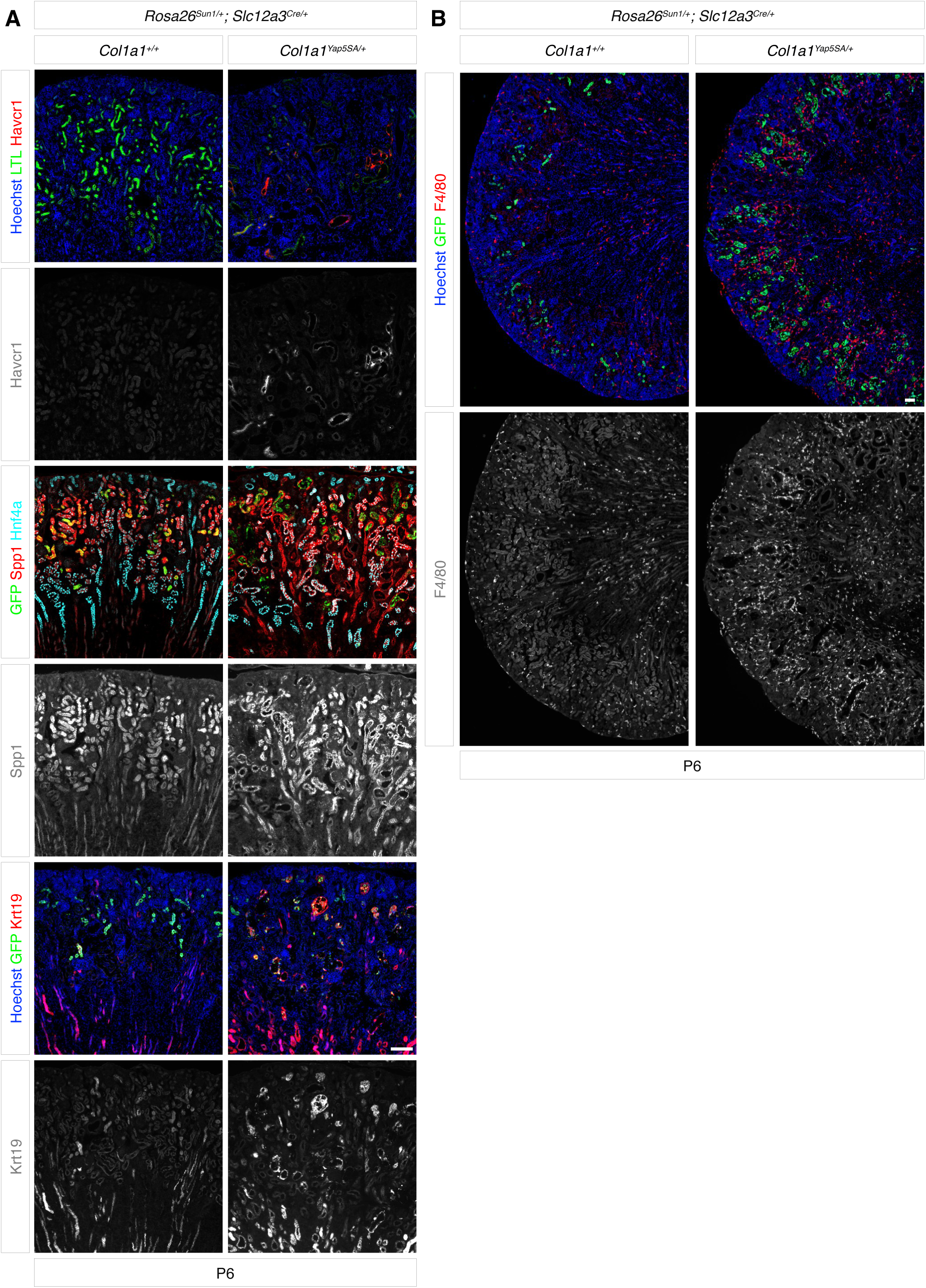
Yap activation in DCT and CNT cells induces tubular injury and macrophage infiltration. (A) In control kidneys, strong LTL labeling is observed with no detectable Havcr1 signal. In contrast, mutant kidneys show reduced LTL labeling accompanied by Havcr1 upregulation. Regions with weak Havcr1 expression overlap with cells displaying weak LTL signal, whereas areas with strong Havcr1 expression show no overlap with LTL. Two additional injury markers, Spp1 and Krt19, are upregulated in mutant kidneys. In control kidneys, Spp1 is largely expressed in Hnf4a+ proximal tubules, whereas in mutant kidneys, Spp1 is upregulated in both Hnf4a+ and Hnf4a-tubules. In control kidneys, Krt19 signal is present in the papilla and absent in GFP+ cells, whereas in mutant kidneys, GFP+ cells show Krt19 upregulation. Stage, P6; Scale bars: 100 µm (B) Mutant kidneys show a pronounced increase in F4/80+ cells, a marker of mature tissue macrophages—reflecting enhanced macrophage infiltration and renal inflammation. Stage, P6; Scale bars: 200 µm

## DISCUSSION

In this study, we investigated the role of the Hippo pathway in distal nephron segments, taking advantage of the fact that constitutive Yap activation functionally mimics Hippo pathway inhibition. To this end, we generated two new mouse lines, a Cre-inducible Yap GOF allele (*Col1a1-Yap5SA*) and the distal nephron-specific *Slc12a3Cre*. These tools allowed selective blockade of Hippo signaling in developing distal nephron epithelia. Constitutive Yap activation produced excessive proliferation and loss of epithelial polarity and junctional organization, disrupted segment identity both within distal nephron segments and within proximal tubules through a non-cell-autonomous mechanism, and caused broad structural disorganization. Together, our findings show that precise Hippo-Yap regulation is essential for maintaining epithelial identity and structural stability across the nephron.

Our results shed new light on the lineage relationship between DCT and CNT segments. Given DCT cells are the only kidney cell type expressing *Slc12a3* in the mouse kidney (2), we initially expected *Slc12a3Cre* to target DCT cells only. Unexpectedly, our lineage tracing analysis showed that *Slc12a3Cre* targets not only Slc12a3-positive DCT cells but also Slc12a3-negative CNT cells (Figure 1A; Supplemental Figure 1A), suggesting that CNT cells emerge from Slc12a3+ DCT cells during development.

Constitutive Yap activation caused DCT and CNT cells to lose expression of segment-specific markers such as *Slc12a3* and *Pvalb* (Figure 1C; Supplemental Figure 1C). These results indicate that appropriate Hippo pathway activity is required to preserve the identity and differentiation state of DCT-derived lineages. Importantly, these findings support a developmental model in which Hippo-Yap signaling is dynamically regulated during nephrogenesis. Prior work has shown that nuclear YAP is present in early nephron epithelia, including the distal domain of the S-shaped body, consistent with a permissive role for YAP during early epithelial expansion (10). Our developmental analyses further indicate that YAP activity becomes progressively repressed in distal nephron epithelia during early postnatal stages, coinciding with terminal differentiation and segmental identity consolidation (Supplemental Figure 3). In this context, sustained YAP activation beyond its normal developmental window disrupts this transition, leading to persistent proliferation, loss of epithelial organization, and impaired differentiation.

Yap functions as a central effector of epithelial proliferation and organ growth, promoting cell-cycle progression and suppressing apoptosis when not restrained by the upstream Hippo kinases MST and LATS (6, 9, 24). Deletion of *Lats1/2* leads to unchecked Yap activation, resulting in excessive proliferation. For example, liver-specific Yap activation causes dramatic organ overgrowth and expansion of progenitor populations (29, 30), demonstrating that loss of Hippo-mediated inhibition drives Yap-dependent hyperplasia. Likewise, *Lats1/2* deletion in adult renal epithelia triggers a rapid, Yap/Taz-dependent proliferative response (16). Consistent with these observations, our data show that constitutive Yap activation in distal nephron epithelia induces marked hyperproliferation (Figure 1B; Supplemental Figure 1B). The morphological and molecular changes in Yap GOF mutant cells closely resemble those characteristics of EMT (31). In these mutant cells, apicobasal polarity was disrupted (Figure 2). The loss of epithelial polarity and junctional integrity likely represents a prerequisite step that enables Yap GOF mutant cells to adopt a mesenchymal-like phenotype. Consistent with this, Yap GOF mutant cells lost their monolayer organization and were detected within neighboring tubular segments, including the loop of Henle and the collecting duct (Figure 3; Supplemental Figure 11). Sustained Yap activity therefore appears sufficient to erode epithelial polarity and promote a less differentiated epithelial state with altered spatial organization within the nephron. Although these findings are consistent with displacement of lineage-labeled distal epithelial cells beyond their normal segmental boundaries, we cannot definitively distinguish active cell migration from alternative explanations based on static analyses alone. While altered segment identity and lineage ambiguity arising from ectopic Slc12a3 expression are considered less likely given the high specificity of Slc12a3 for the distal convoluted tubule and the absence of *Slc12a3Cre*-mediated recombination in adjacent segments under normal conditions, resolving these possibilities conclusively will require future studies employing live imaging or single-cell resolution temporal lineage tracing.

In addition to structural disruption, constitutive Yap activation in distal nephron epithelia induced a strong inflammatory response. RNA-seq analysis revealed marked upregulation of TNF, MAPK, and NF-κB pathway genes, indicating activation of inflammatory and immune signaling (Supplemental Figure 15B). Consistent with this, immunostaining showed extensive macrophage infiltration and activation (Figure 6B). The mutant kidneys also exhibited strong induction of injury-associated markers, including *Havcr1* and *Spp1* (Figure 6A). *Havcr1* (Kim1) is transiently expressed during repair but becomes maladaptive when chronically elevated (32, 33). *Spp1* (Osteopontin) functions as a proinflammatory factor that promotes apoptosis and fibrosis during kidney injury (34, 35). Together, these results indicate that sustained Yap activation in distal nephron segments elicits chronic injury responses characterized by persistent inflammatory signaling, macrophage recruitment, and maladaptive remodeling.

*YAP5SA* expression in distal nephron segments (DCT/CNT) impaired proximal tubule differentiation through a non-cell-autonomous mechanism. Both bulk RNA-seq and immunofluorescence analyses supported this conclusion. Although Hnf4a+ proximal tubule cells were present in the mutant kidneys, they showed reduced or absent expression of several proximal tubule markers (Figure 5), indicating that proximal tubule specification occurs but maturation is arrested when Yap is constitutively activated in distal nephron segments.

Developmental time⍰course analyses at E18.5 and P3 demonstrated that proximal tubule identity defects progressively emerge after distal Yap activation (Supplemental Figure 16). Importantly, these early phenotypes arise before overt tissue degeneration or terminal pathology, supporting the conclusion that transcriptional alterations detected at P3 largely reflect direct consequences of sustained YAP activation rather than nonspecific degeneration.

Interpretation of the temporal onset and penetrance of these phenotypes must be considered in the context of the asynchronous nature of nephrogenesis. In the mouse kidney, nephrons are specified in successive waves rather than synchronously, such that individual distal tubule segments acquire Cre-mediated recombination and sustained YAP activation at different absolute times. As a result, distal nephron cells across different nephrons experience variable durations of aberrant YAP activity at any given stage, explaining the heterogeneous defects observed at E18.5 and the more uniform phenotypes seen by P3 and P6.

The mechanism underlying this distal-to-proximal effect remains unclear, but we propose three non-mutually exclusive possibilities. First, distal nephron epithelia may normally produce paracrine cues required for proximal tubule maturation. Second, strong inflammatory activation seen in Yap GOF kidneys may destabilize proximal tubule identity. Third, the structural disorganization and polarity loss in Yap GOF DCT and CNT segments may compromise epithelial integrity and impair luminal flow. Loss of flow would deprive proximal tubule cells of shear-stress signals necessary for proper polarization and maturation (28). Flow-dependent morphogenesis has been shown in zebrafish nephrons and in human organoid proximal tubules, where flow promotes transporter expression and maturation (36, 37). Future work will be required to determine the relative contribution of each mechanism.

Consistent with a flow⍰based model, we observed disruption of the MD-JGA axis in Yap GOF kidneys. Constitutive Yap activation in distal nephron segments resulted in loss of Renin expression and coordinated downregulation of multiple renin-angiotensin system (RAS) components, including *Ren1*, *Ace*, *Agtr1a*, and *Agtr2* (Supplemental Figure 12; Supplemental Table 1). Functional assessment using an in vivo fluorescent dextran delivery assay revealed markedly reduced filtration⍰dependent luminal delivery in mutant kidneys, providing direct evidence of impaired effective nephron flow (Supplemental Figure 13). Together, these findings support the idea that distal epithelial disorganization can compromise upstream filtration and downstream tubular delivery, thereby contributing to non⍰ce⍰llautonomous defects in proximal tubule maturation.

These alterations recapitulate key features of Renal Tubular Dysgenesis (RTD), a congenital disorder caused by autosomal recessive mutations in the renin-angiotensin system genes (e.g., *REN*, *AGT*, *ACE*, *AGTR1*) and characterized by severely underdeveloped proximal tubules, oligohydramnios, profound hypotension, and perinatal lethality (38). Although mouse models of RTD remain limited (39), our Yap GOF mutant mouse model shows reduced Renin with proximal tubule maturation failure, suggesting that dysregulated Hippo-Yap signaling in the distal nephron can phenocopy core molecular and developmental features of this condition.

In summary, our findings identify Hippo-Yap balance in DCT/CNT as a critical regulator of nephron patterning and inter-segmental communication. Constitutive Yap activation drives hyperproliferation, epithelial architectural destabilization, inflammatory reprogramming, and a non-cell-autonomous failure of proximal tubule maturation linked to impaired MD-JGA signaling and nephron flow. These results argue that Yap activity must be precisely constrained during development and reveal a previously unrecognized level of functional interdependence among nephron segments. More broadly, our study provides a framework for understanding how perturbations within one tubular compartment can propagate across the nephron to disrupt global epithelial organization and endocrine-physiological integration.

## METHODS

### Sex as a biological variable

Both male and female mice were used in this study, with equal representation of each sex. Mutant and control animals of both sexes were included in all analyses unless otherwise indicated. Sex-based differences were assessed, and no appreciable differences were observed between male and female mice across the measured outcomes. Therefore, data from male and female animals were pooled for subsequent analyses.

### Generation of Slc12a3-IRES-Cre

The *Slc12a3-IRES-Cre* mouse line was generated by CRISPR/Cas9-mediated modification of *Slc12a3-IRES-CreERT2* (JAX: 030602) (20). Briefly, the Cas9 protein (#1081061, IDT) was incubated with the single-guide RNA (GGCGATCTCGAGCCATCTGC) before microinjection into fertilized embryos. The donor oligo designed to insert a stop codon at the end of the coding sequence of Cre (40) was also added to form the ribonucleoprotein (RNP) complex (41). The final concentrations were 60 ng/ml single-guide RNA, 80 ng/ml Cas9 protein, and 500 ng/ml donor oligo. Zygotes carrying *Slc12a3-IRES-CreERT2* were collected from superovulated females and electroporated on ice with 7 μl of the RNP/donor mix using a Genome Editor electroporator (BEX). Zygotes were transferred into 500 μl of M2 medium (Sigma), and transplanted into the oviductal ampullae of pseudopregnant CD-1 females. Offspring were genotyped by PCR, EcoRI digestion, and Sanger sequencing to confirm successful genome editing. Genotyping PCR was performed using three oligonucleotides (AGA GGG TGC GTT CTG ACT CT, GTC CCA GGA CCT CAG ACC TC, and TAC GCT TGA GGA GAG CCA TT). This primer set yields a 150 bp amplicon for the wild-type *Slc12a3* allele and a 500 bp amplicon for the *Slc12a3-IRES-Cre* allele.

### Generation of a Cre-inducible *Yap* gain-of-function allele (*Col1a1-Yap5SA*)

To create a mouse model enabling conditional expression of a constitutively active Yap variant (*Yap5SA*), we employed a CRISPR/Cas9-mediated knock-in strategy targeting the *Col1a1* safe harbor locus (42, 43). In this system, Col1a1 was used solely as a genomic docking site for targeted integration and was not used as a promoter for Yap expression. The *Yap5SA* variant carries five serine-to-alanine mutations (S61A, S109A, S127A, S164A, S381A), rendering it resistant to LATS kinase-mediated inhibition and resulting in constitutive activation (22, 23). To decouple transgene expression from endogenous Col1a1 regulation, the constitutively active Yap5SA cassette was placed under the control of the constitutive CAG promoter, ensuring Yap expression independent of ce⍰lltype-specific Col1a1 activity. A donor vector contains a CAG promoter, lox66, an inverted 3xFLAG-tagged *Yap5SA* followed by an SV40 polyadenylation signal, lox71, and two copies of insulators (44) (Supplemental Figure 4). Heterotypic lox66 and lox71 sites enable Cre-dependent unidirectional inversion (45). This cassette was cloned in reverse orientation relative to the CAG promoter within the targeting vector, which also included 5′ and 3′ homology arms of 2.5 kb and 3.0 kb, respectively, to promote homologous recombination. The final construct was amplified and purified using the EndoFree Plasmid Kit (Qiagen). Pronuclear injections were performed on fertilized C57BL/6J zygotes with a mixture containing Cas9 protein at 40 ng/μL, chemically modified single-guide RNA (GGGAGGAAACCTGCCCTTGG) at 30 ng/μL, and the donor vector at 6 ng/μL. Injected embryos were transferred into the oviductal ampullae of pseudopregnant CD-1 females (approximately 25 embryos per recipient). Resulting pups were screened for correct integration via long-range PCR and confirmed by Sanger sequencing. Genotyping PCR was performed using three primers: CTGAACTCAGCATCCGAGCTGTAC, ACTCCATATATGGGCTATGAACTAATGACC, and CAGCCCTGCACAGAATTGTCAG. This reaction produces a 344 bp amplicon for the wild-type *Col1a1* allele and a 285 bp amplicon for the *Col1a1-Yap5SA* allele.

### Mice and breeding strategy

*Rosa26-Sun1* (JAX: 021039) expresses a nuclear membrane protein SUN1 fused to superfolder GFP upon Cre-mediated recombination (21). *Rosa26-NuTRAP* (JAX: 029899) expresses the 60S ribosomal subunit L10a fused to EGFP upon Cre-mediated recombination (46). For genetic experiments, *Slc12a3^Cre/Cre^* mice were crossed with *Col1a1^Yap5SA/+^; Rosa26^Sun1/Sun1^* mice to generate *Slc12a3^Cre/+^; Col1a1^Yap5SA/+^; Rosa26^Sun1/+^*mutant mice. Littermate *Slc12a3^Cre/+^; Col1a1^+/+^; Rosa26^Sun1/+^* mice were used as controls. In a subset of experiments presented in the Supplemental Figures, *Rosa26-NuTRAP* was used instead of *Rosa26-Sun1*. All mice in this report were bred and maintained in a mixed genetic background. Animals were centrally managed by Northwestern University Center for Comparative Medicine (CCM), an animal care facility that operates within full compliance of the NIH guidelines. Mice were monitored daily and housed in a controlled environment with a 12-hour light/12-hour dark cycle, with ad libitum access to water and a standard chow diet. At the first sign of pain, suffering or distress, mice were euthanized. Humane end points were established to include lethargy, labored breathing, or swelling.

### Bulk RNA-seq

Kidneys were collected at postnatal day 3 from two mutants and two controls of each sex. Total RNA was extracted from postnatal day 3 kidneys using the Qiagen RNeasy Plus Micro Kit. mRNA was isolated from total RNA using the NEBNext Poly(A) mRNA Magnetic Isolation Module (E7490L, New England Biolabs). Fragmentation of mRNA and cDNA synthesis were performed using NEBNext RNA First Strand Synthesis Module (E7525L) and NEBNext RNA Second Strand Synthesis Module (E6111L). The cDNA’s were processed to sequencing libraries using ThruPLEX DNA-seq 12S Kit (R400675 and R400695, Takara). Libraries were sequenced on Illumina NovaSeq X Plus or Element AVITI at the NUSeq Core facility at Northwestern University.

### RNA-seq analysis

Paired-end RNA-seq reads were mapped to the UCSC mouse reference genome (mm10) using the STAR aligner (47). Only uniquely aligned reads were used for differential gene analysis. Gene-level read counts were obtained using the FeatureCounts tool from the Subread package, with the following parameters: “-s 2 -O --fracOverlap 0.8” (48). Differential expression analysis was conducted using the DESeq2 package (49). Genes exhibiting a fold-change > 1.5 and FDR < 0.05 were considered significantly differentially expressed. Gene expression data for all genes are provided in Supplemental Table 1. Gene ontology (GO) analysis was performed using EnrichR (50). Gene expression of select genes were visualized as dot plots in single-cell RNA-seq datasets from mouse kidneys at embryonic day 18.5 and postnatal day 0 (GSE214024 and GSE275601) (40) using Seurat (51). Module scores of the differentially expressed genes from the bulk RNA-seq data were calculated using the AddModuleScore function in Seurat and projected to UMAP.

### Immunofluorescence staining and microscopy

Kidneys from mutant and control animals of both sexes were analyzed. For immunofluorescence analyses, biological replicates consisted of kidneys collected from independent animals; two control and four mutant biological replicates were examined, unless otherwise indicated. Tissue preparation, staining and imaging were performed in parallel for each experiment to minimize technical variation. Sample size was determined by litter size and breeding outcomes. No formal sample size calculation was performed. No exclusion criteria were defined, and no animals or data points were removed from the analysis. Samples were processed in randomized order, and control and mutant kidneys were embedded within the same block to reduce batch effects. The investigator (Z.D.G.) was aware of sample genotypes during processing and analysis. Kidneys were fixed in phosphate-buffered saline (PBS) containing 4% paraformaldehyde (PFA) for 20 minutes and then incubated overnight at 4°C in PBS containing 10% sucrose. Tissues were embedded in OCT compound (Thermo Fisher Scientific) and stored at −80°C. Cryosections (8 μm) were incubated overnight at 4°C in PBS containing 0.1% Triton X-100, 5% heat-inactivated sheep serum, and primary antibodies (Supplemental Table 2).

Fluorophore-labeled secondary antibodies were sourced from Invitrogen or Jackson ImmunoResearch (Supplemental Table 2). Images shown in Figure 2A were acquired at the Center for Advanced Microscopy/Nikon Imaging Center (RRID:SCR_020996) at Northwestern University using a Nikon CSU⍰W1 SoRa super⍰resolution spinning disk confocal system, which was purchased with support from NIH S10 award 1S10OD032270⍰01A1. All other images were obtained using a Nikon Ti2 widefield microscope.

### Cell proliferation quantification

Cell proliferation was assessed by immunofluorescent staining for Ki67. Approximately three non-overlapping fields per kidney (field size: 389.6 µm × 389.6 µm) were analyzed across three biological replicates. GFP+ cells and GFP+/Ki67+ double-positive cells were manually counted using ImageJ/Fiji. The proliferation index was calculated as the percentage of GFP+/Ki67+ cells relative to the total number of GFP+ cells. Field-level measurements were averaged within each biological replicate prior to statistical analysis.

### In vivo fluorescent dextran delivery assay

Postnatal day 3 pups of both sexes were injected retro-orbitally with 5 µl of 10-kDa Texas Red-conjugated dextran (5 mg/ml; Fisher Scientific, D1863), as described previously (28). Pups were returned to the dam and euthanized after a 5-minute chase period. Kidneys were processed for immunofluorescence staining using the same protocol as described above. Uninjected littermates were processed in parallel as negative controls. Experiments were performed using three independent biological replicates, including both male and female pups.

### Histology

Kidneys were fixed in 4% PFA in PBS overnight and submitted to the Mouse Histology & Phenotyping Laboratory (MHPL) at the Robert H. Lurie Comprehensive Cancer Center of Northwestern University which is supported by the National Cancer Institute (NCI) P30-CA060553. Paraffin sections (3 μm) were stained with hematoxylin and eosin at MHPL, and images were taken with a Nikon Ti2 widefield microscope housed at the Center for Advanced Microscopy/Nikon Imaging Center (RRID:SCR_020996) at Northwestern University.

## Supporting information

Supporting data values

## Statistical analyses

All statistical analyses were performed using GraphPad Prism version 10. Kaplan-Meier survival curves were generated and analyzed using the log-rank (Mantel-Cox) test (mutant mice, n = 51; control mice, n = 51). For all other quantitative analyses, comparisons between two groups were performed using unpaired two-tailed Student’s t-tests in GraphPad Prism. No statistical methods were used to predetermine sample size, and data distribution was not formally tested for normality. A P value < 0.05 was considered statistically significant.

## Study approval

All animal experiments were approved by the Institutional Animal Care and Use Committees (IACUC) at Northwestern University and Cincinnati Children’s Hospital Medical Center and were conducted in accordance with the NIH Guide for the Care and Use of Laboratory Animals.

## Data availability

The bulk RNA sequencing data generated in this study have been deposited in the Gene Expression Omnibus (GEO) under accession number GSE297246. All other data supporting the findings of this study are available from the corresponding author upon reasonable request. Source data for all graphs are provided in the Supporting Data Values Excel file.

## Author Contributions

ZDG and JSP designed the study. ZDG, EC, HAC, AA, and JSP conducted experiments. MS, CA, and HWL performed bioinformatic data analysis. ZDG and JSP curated and validated the data and prepared the figures. BRT and YCH contributed to methodology development. JSP acquired funding, provided resources, and administered the project. ZDG and JSP wrote the original draft of the manuscript, and ZDG, EC, and JSP reviewed and edited the final version. All authors approved the final version of the manuscript.

## Supplemental Material

Supplemental Figure 1. Constitutive activation of Yap in the distal convoluted tubule (DCT) and connecting tubule (CNT) leads to increased proliferation and disrupt segmental identity

Supplemental Figure 2. Schematic representation of ion transport, cell types, and transport proteins in the distal nephron and collecting duct

Supplemental Figure 3. Developmental downregulation of nuclear Yap in distal nephron segments.

Supplemental Figure 4. Cre-dependent activation of constitutively active Yap5SA

Supplemental Figure 5. Specific Expression of the FLAG-tagged Yap transgene in distal nephron segments

Supplemental Figure 6. Constitutive activation of Yap impairs distal nephron segment identity at E18.5

Supplemental Figure 7. Constitutive activation of Yap impairs distal nephron segment identity at P3

Supplemental Figure 8. Disrupted nephron structure and postnatal lethality caused by Yap activation in the distal nephron segments

Supplemental Figure 9. Constitutive activation of Yap in DCT and CNT leads to progressive loss of epithelial polarity

Supplemental Figure 10. Constitutive activation of Yap in DCT and CNT impairs epithelial polarity and junctional organization

Supplemental Figure 11. Constitutive Yap activation in DCT and CNT leads to aberrant localization of lineage⍰labeled cells within the collecting duct.

Supplemental Figure 12. Constitutive Yap activation in distal nephron segments disrupts macula densa integrity and renin expression

Supplemental Figure 13. Constitutive activation of Yap in distal nephron tubules reduces luminal dextran accumulation

Supplemental Figure 14. Integrating Bulk RNA-seq differential expression with single-nucleus RNA-seq to identify cell-type-specific responses to Yap activation

Supplemental Figure 15. Gene Ontology and KEGG pathway enrichment analyses of differentially expressed genes in Yap GOF kidneys

Supplemental Figure 16. Constitutive activation of Yap in DCT and CNT affects proximal tubule development

Supplemental Figure 17. Constitutive Yap activation in distal nephron segments perturbs proper papilla formation

Supplemental Table 1. RNA-seq analysis comparing Yap GOF mutant kidneys and controls at postnatal day 3

Supplemental Table 2. Primary and secondary antibodies used in this study

