## Supplemental Figures 1-17 for "Constitutive Yap activation in distal nephron segments disrupts epithelial identity and nephron patterning"

### Supplemental Figure 1

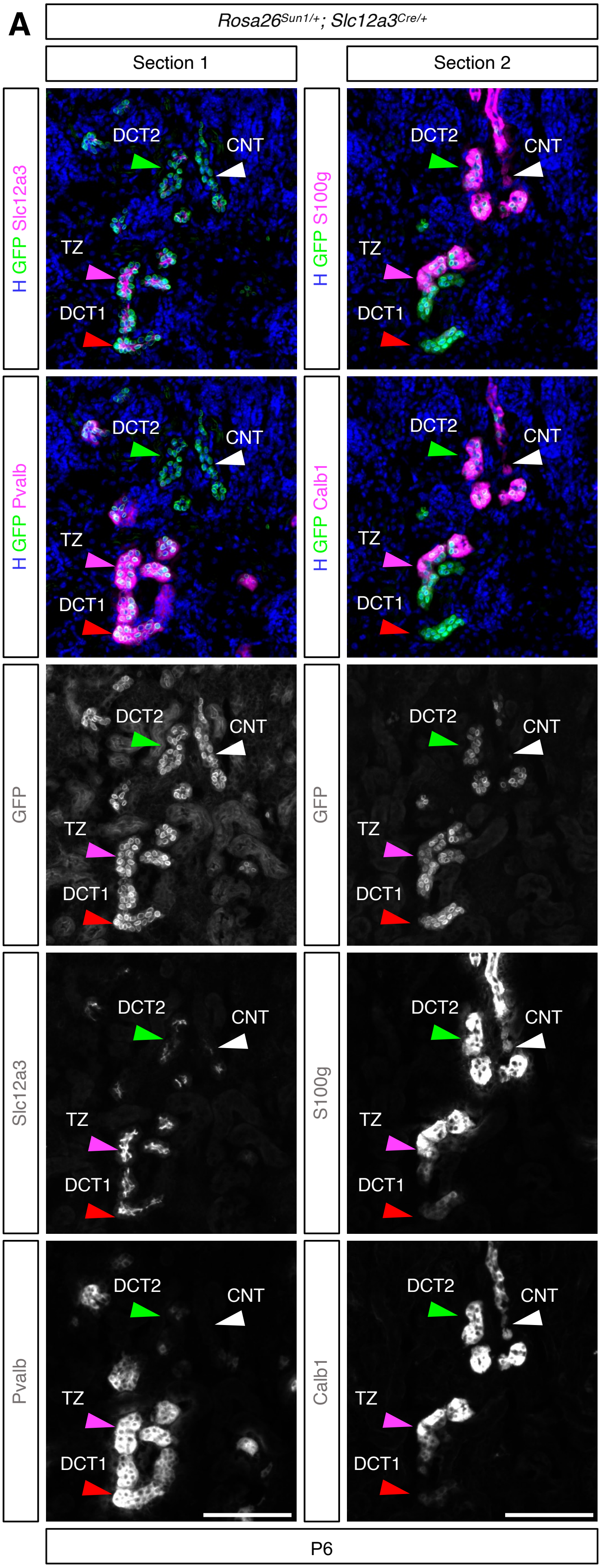

### Supplemental Figure 1

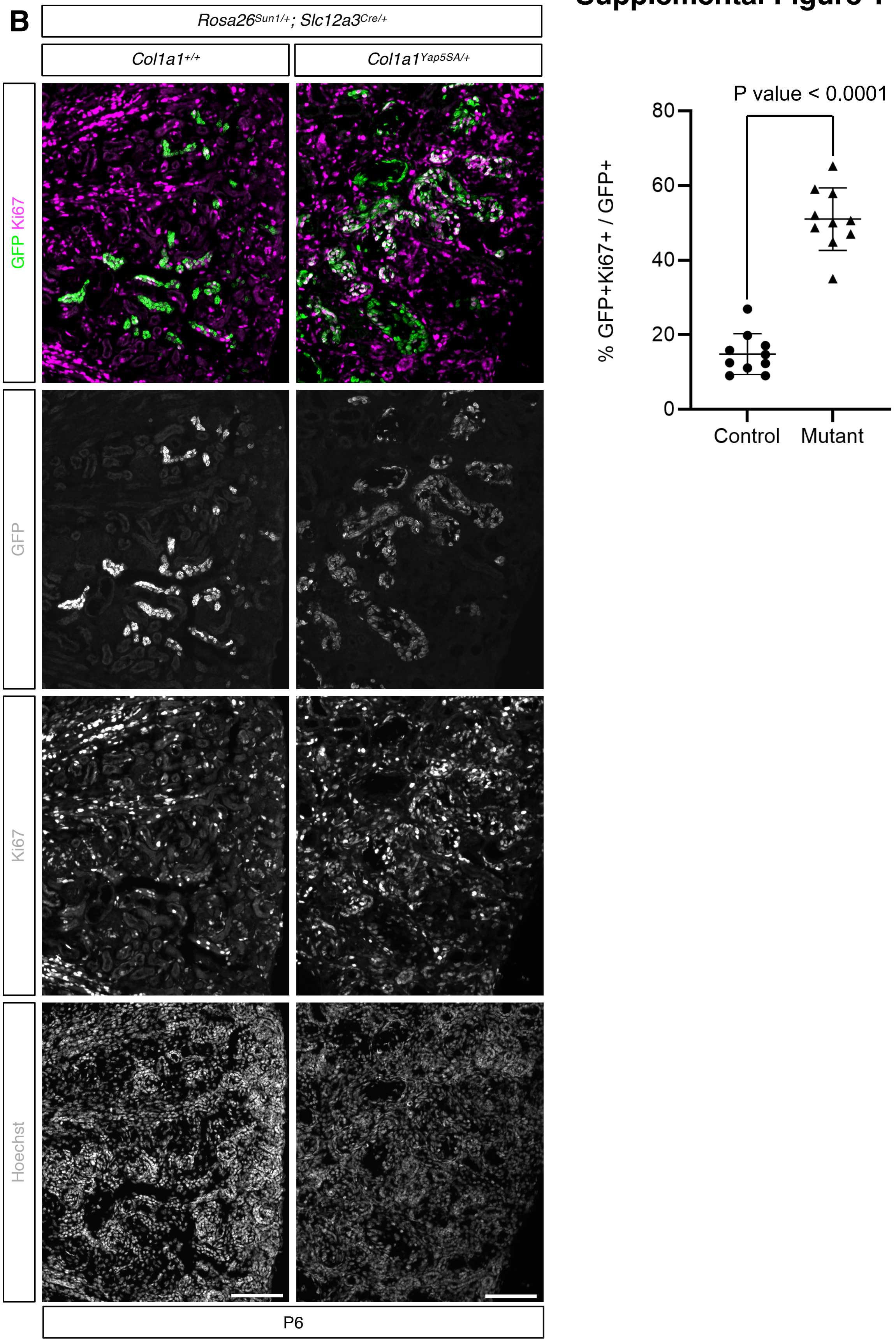

### Supplemental Figure 1

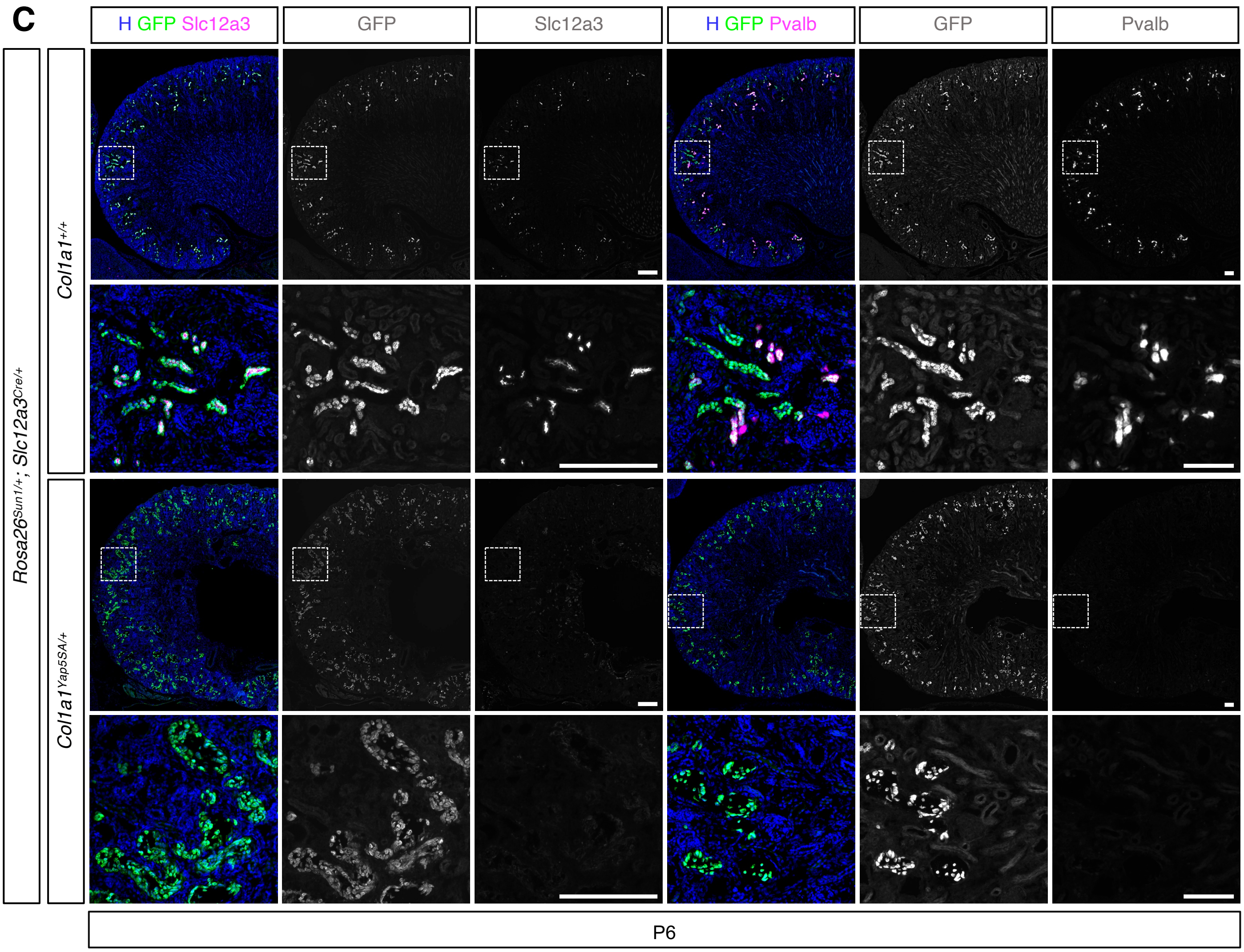

### Supplemental Figure 1

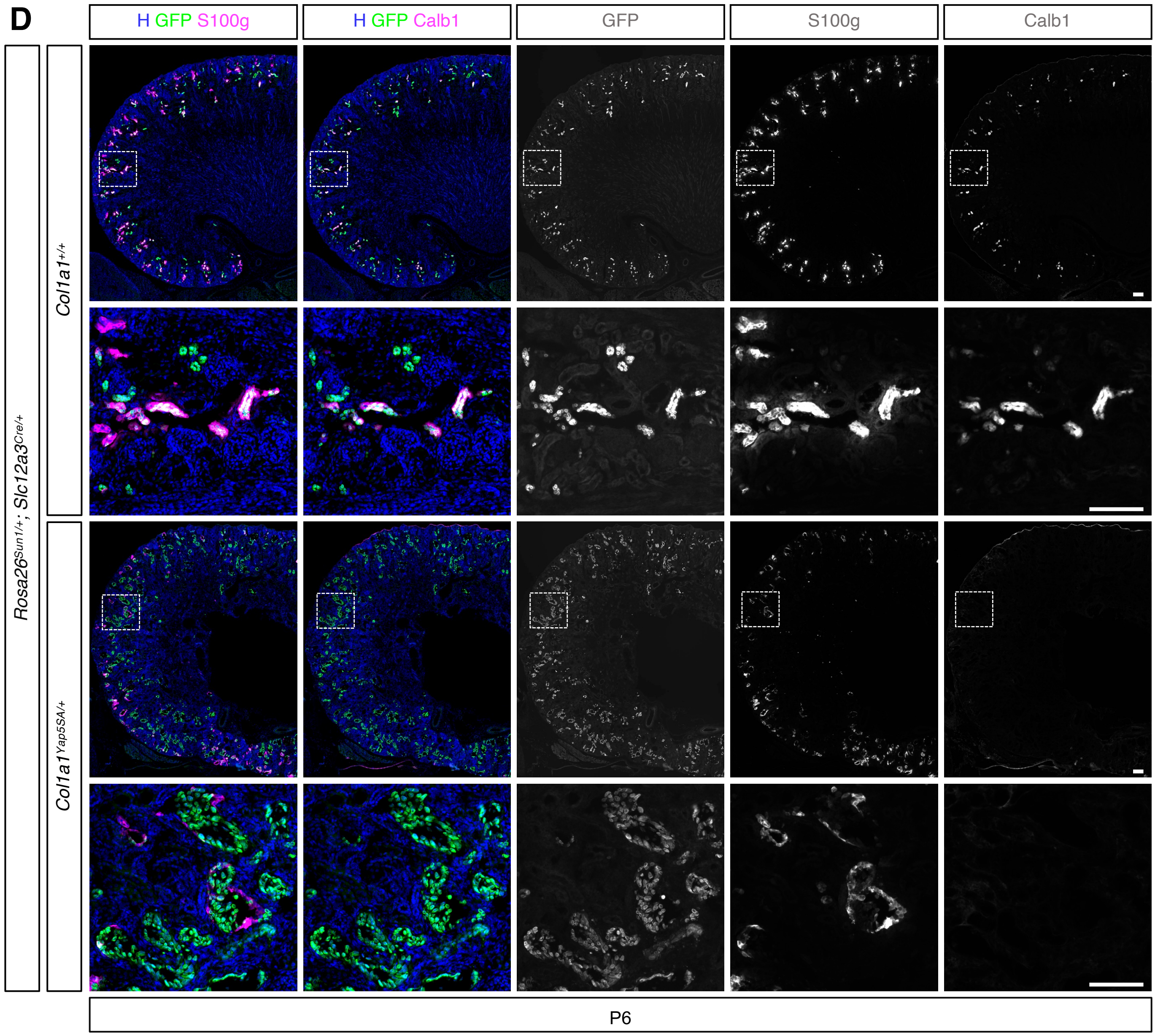


**Supplemental Figure 1. Constitutive activation of Yap in the distal convoluted tubule (DCT) and connecting tubule (CNT) leads to increased proliferation and disrupt segmental identity.** Data are identical to Figure 1; colors were adjusted, and single-channel grayscale images were included to improve accessibility for readers with color-vision deficiencies. (A) Lineage tracing using the *Rosa26-Sun1* reporter (nuclear membrane GFP) shows that *Slc12a3Cre* targets both DCT and CNT. Slc12a3 marks both DCT1 and DCT2, while Pvalb marks DCT1 only. S100g and Calb1 mark DCT2 and CNT. The transition zone (TZ) is marked by both DCT1 and DCT2 markers (see Supplemental Figure 2). Adjacent sections of the same kidney are shown. (B) In control kidneys, only a small subset of DCT and CNT cells (GFP+) are positive for Ki67, indicating low proliferative activity. In contrast, DCT and CNT cells with persistent Yap activation show strong Ki67 staining, consistent with increased proliferation. Quantification of Ki67+ cells among GFP+ distal nephron cells demonstrates a significant increase in mutants compared with controls ($P < 0.0001$). Data are presented as mean ± SEM. (C, D) GFP+ cells in the control kidney express markers characteristic of distal nephron segments, while GFP+ cells in the mutant kidney lack these markers, suggesting disrupted segmental identity. (H) Hoechst labels nuclei. (A-D) Stage: postnatal day 6; Scale bar: 100 $\mu$m.

### Supplemental Figure 2

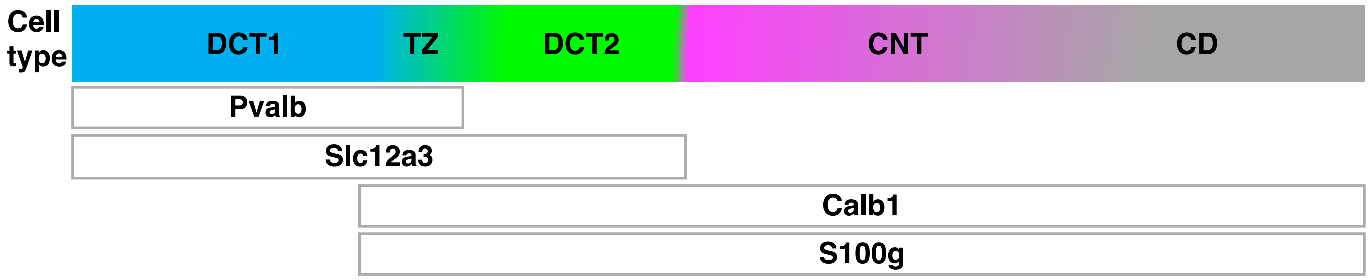


**Supplemental Figure 2. Schematic representation of ion transport, cell types, and transport proteins in the distal nephron and collecting duct**

DCT1 refers to the early distal convoluted tubule, DCT2 to the late distal convoluted tubule, CNT to the connecting tubule, and CD to the collecting duct. Slc12a3 labels both DCT1 and DCT2, Pvalb labels DCT1, and S100g and Calb1 label DCT2 and CNT. The transition zone (TZ) contains cells co-expressing DCT1 and DCT2 markers.

### Supplemental Figure 3

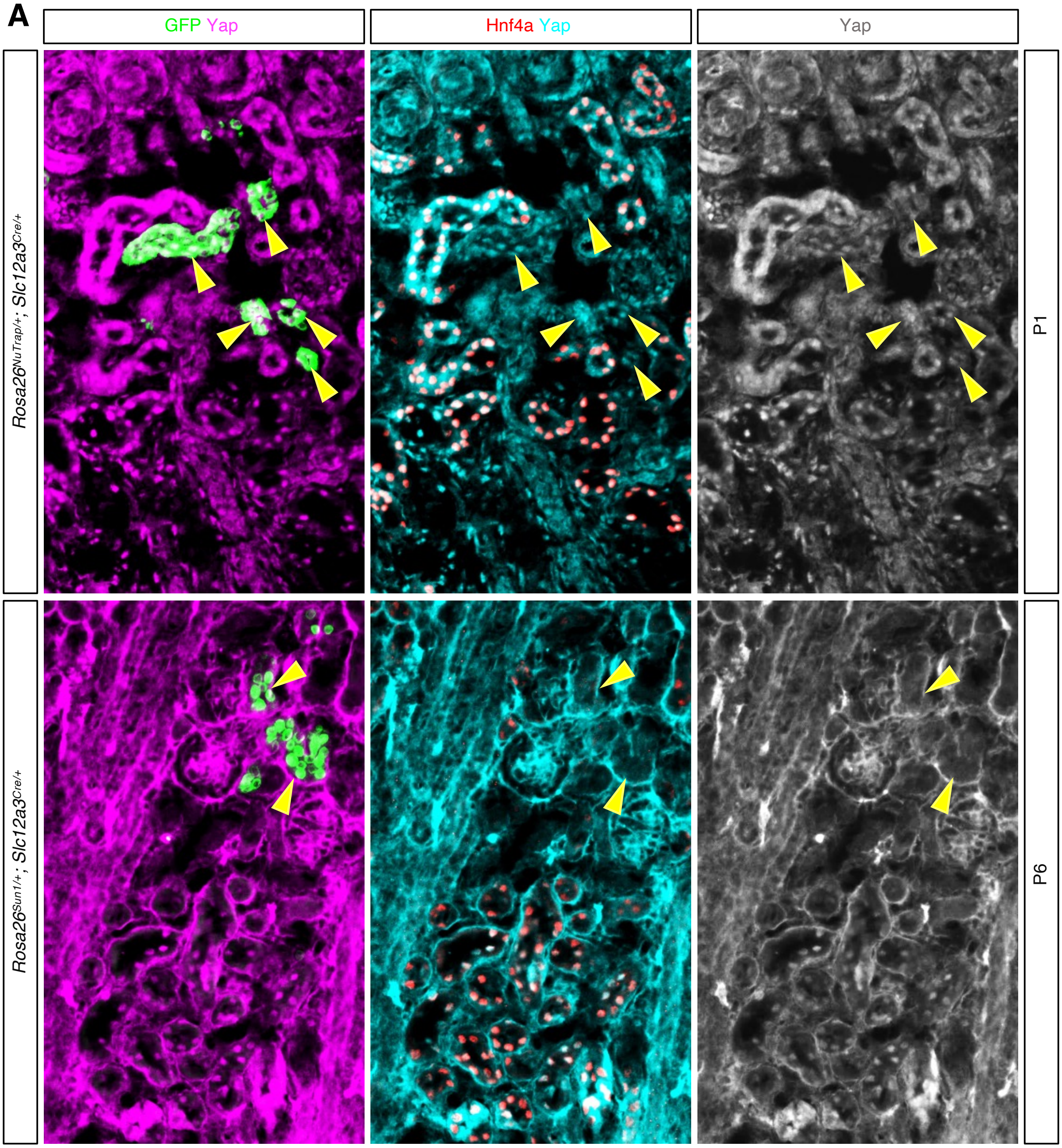

**B**

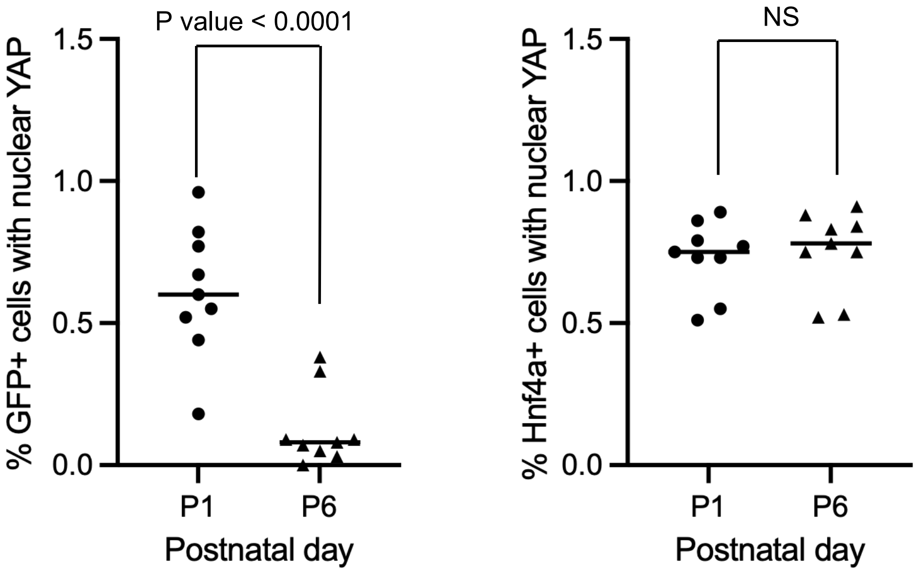


**Supplemental Figure 3. Developmental downregulation of nuclear Yap in distal nephron segments.** (A) At postnatal day 1 (P1), Yap is prominently localized to the nuclei of both proximal tubule cells (Hnf4a+) and distal nephron segments (GFP+) (arrowheads). By postnatal day 6 (P6), nuclear Yap remains evident in proximal tubule cells but is markedly reduced in distal nephron segments (arrowheads), indicating a developmental downregulation of Yap activity during postnatal maturation of the distal nephron. Individual fluorescence channels are shown in grayscale for clarity. (B) Quantification of nuclear Yap signal shows a significant reduction in GFP+ distal nephron cells at P6 compared to P1 (***$P < 0.001$), whereas nuclear Yap levels in Hnf4a+ proximal tubule cells are unchanged (not significant). Scale bar: 50 $\mu$m.

### Supplemental Figure 4

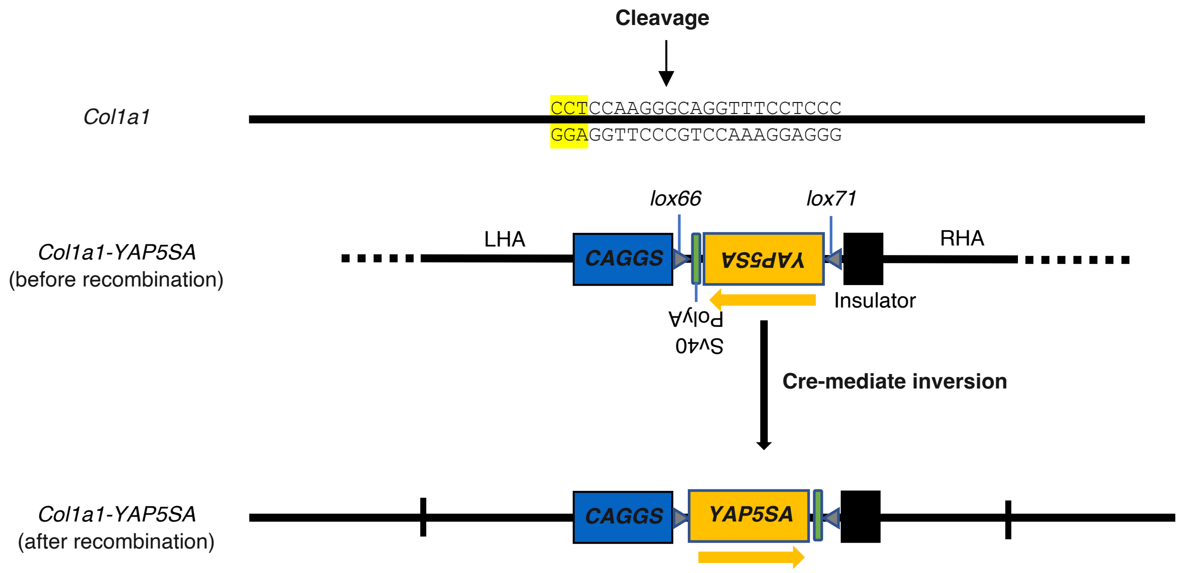


**Supplemental Figure 4. Cre-dependent activation of constitutively active Yap5SA**
A donor cassette containing inverted *YAP5SA* (S61A/S109A/S127A/S164A/S381A) is flanked by left and right homology arms (LHA and RHA) and mutant lox sites (*lox66* and *lox71*). This cassette was inserted into the *Col1a1* locus by CRISPR–Cas9, targeting a site upstream of a CCT–GGA protospacer-adjacent motif (PAM, highlighted in yellow). Upon Cre expression, the lox66/lox71 pair undergoes a single, irreversible recombination event that inverts the cassette into the sense orientation (orange arrow), thereby enabling CAGGS-driven expression of *YAP5SA*.

### Supplemental Figure 5

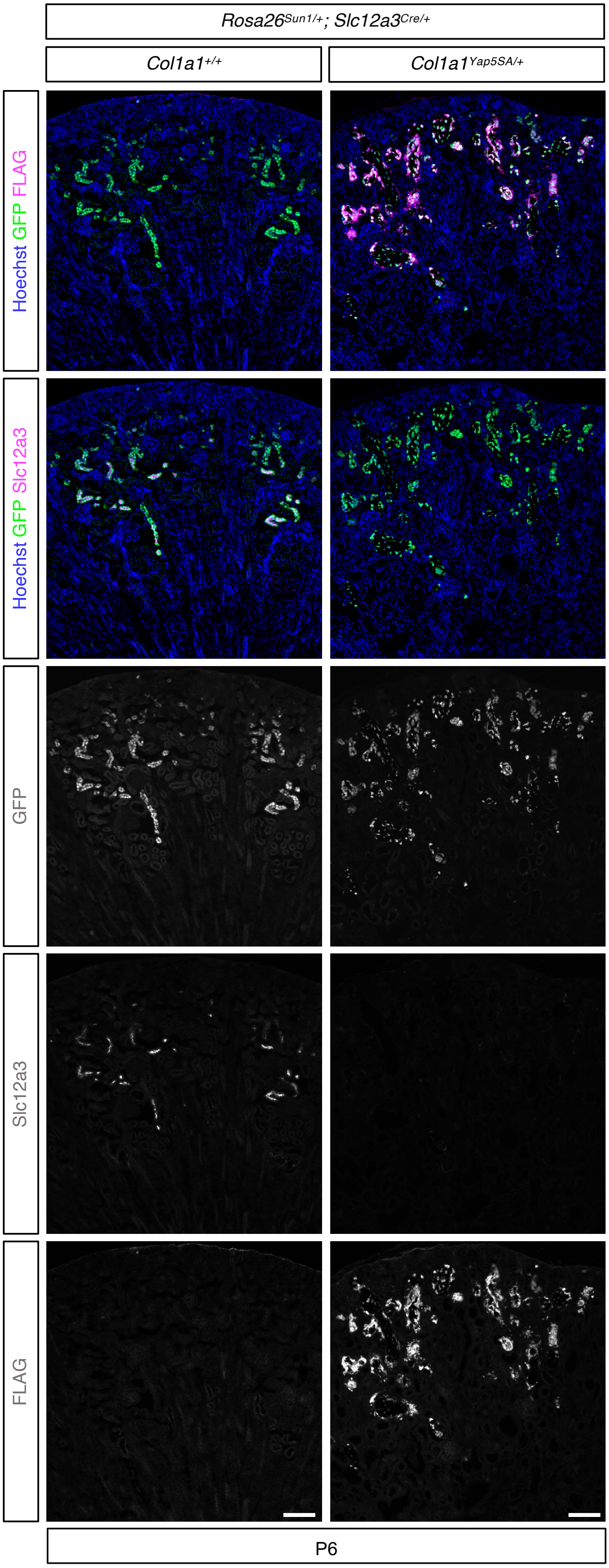


**Supplemental Figure 5. Specific Expression of the FLAG-tagged Yap transgene in distal nephron segments.** In mutant kidneys, all GFP+ distal nephron epithelial cells are FLAG+, confirming expression of the 3xFLAG-tagged Yap transgene in the targeted population. GFP+/FLAG+ cells lack Slc12a3 expression, indicating loss of distal convoluted tubule (DCT) identity following constitutive Yap activation. Stage: postnatal day 6; Scale bar: 100 $\mu$m.

### Supplemental Figure 6

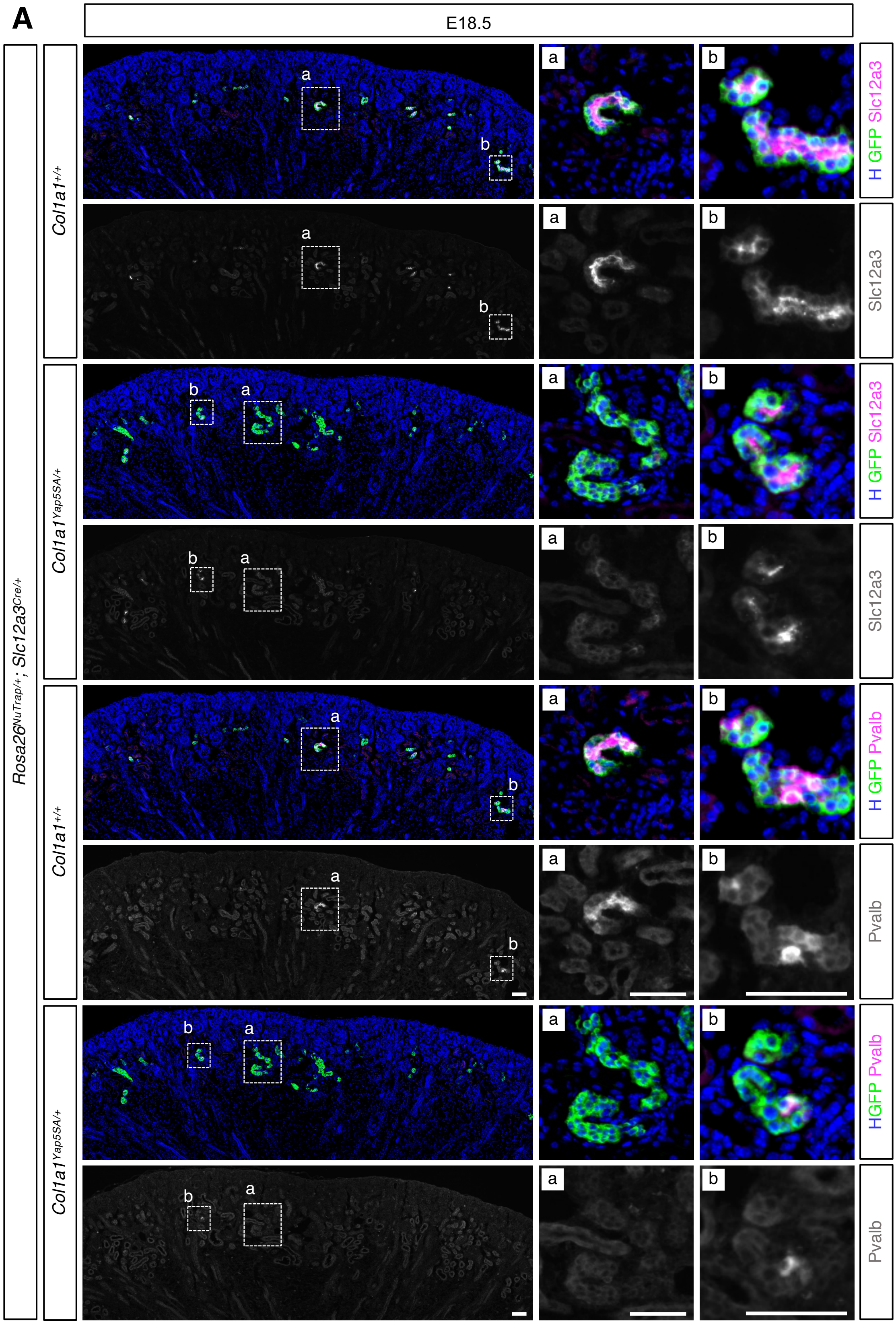

### Supplemental Figure 6

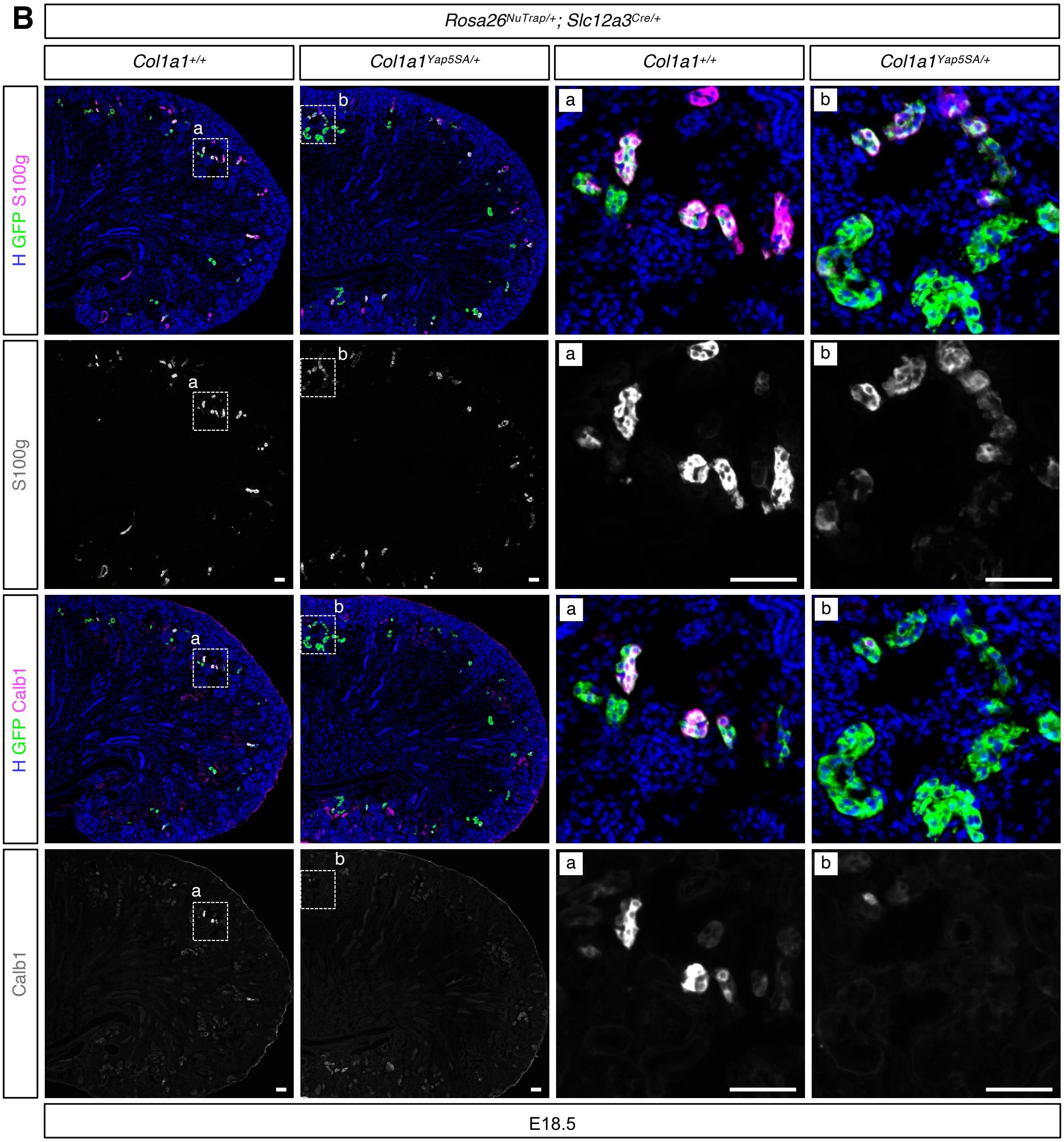


**Supplemental Figure 6. Constitutive activation of Yap impairs distal nephron segment identity at E18.5.** At embryonic day 18.5 (E18.5), GFP+ cells in control kidneys express distal nephron markers, including Slc12a3 and Pvalb (A) and S100g and Calb1 (B). In mutant kidneys, GFP+ cells show heterogeneous marker expression, with some tubules lacking these markers while others retain expression. The left panels show low-magnification views of the kidney, and the regions outlined by dotted squares are shown at higher magnification in the right panels. In (A), separate insets show GFP+ tubules lacking (a) or retaining (b) Slc12a3 and Pvalb expression in mutant kidneys. In (B), a single higher-magnification inset of a mutant kidney shows both marker-negative and marker-positive GFP+ tubules for S100g and Calb1 within the same field. Individual fluorescence channels are shown in grayscale for clarity. Scale bar: 50 $\mu$m.

### Supplemental Figure 7

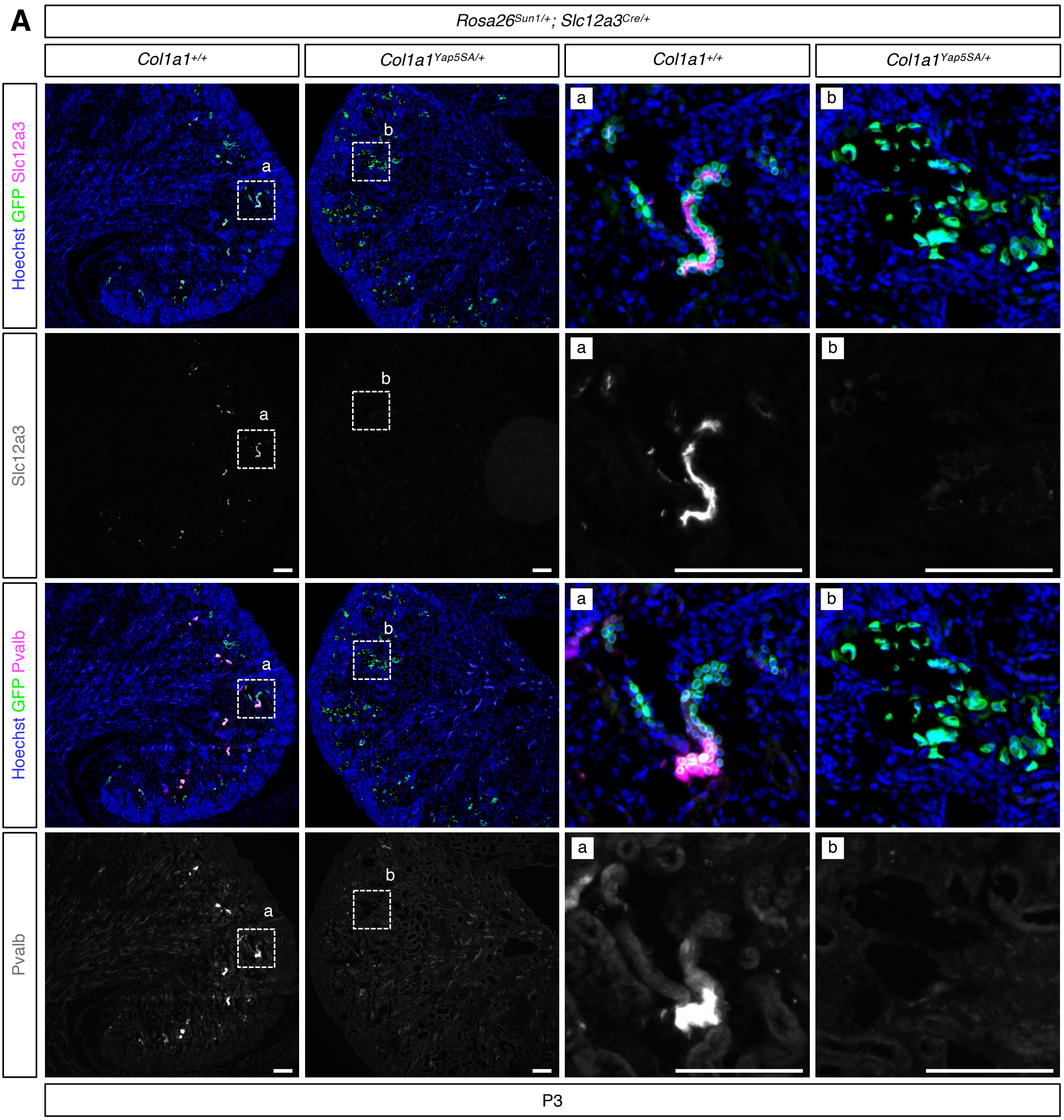

### Supplemental Figure 7

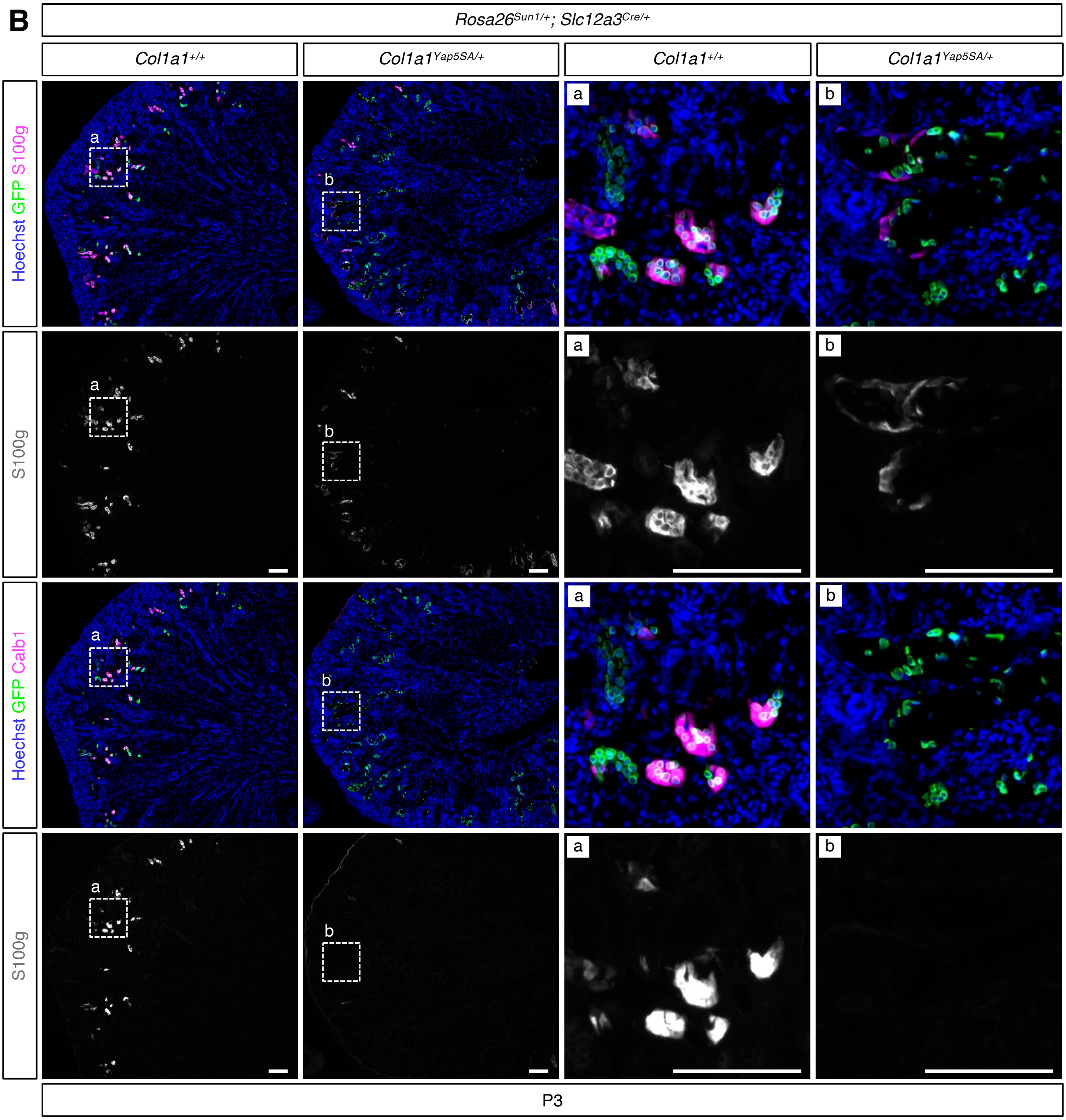


**Supplemental Figure 7. Constitutive activation of Yap impairs distal nephron segment identity at P3.** At postnatal day 3 (P3), GFP+ cells in control kidneys express distal nephron markers, including Slc12a3 and Pvalb (A) and S100g and Calb1 (B). In mutant kidneys, nearly all GFP+ cells lack expression of these markers, indicating widespread loss of distal convoluted tubule (DCT) and connecting tubule (CNT) identity. The left panels show low-magnification views of the kidney, with boxed regions shown at higher magnification on the right. Individual fluorescence channels are presented in grayscale for clarity. Scale bar: 100 $\mu$m.

### Supplemental Figure 8

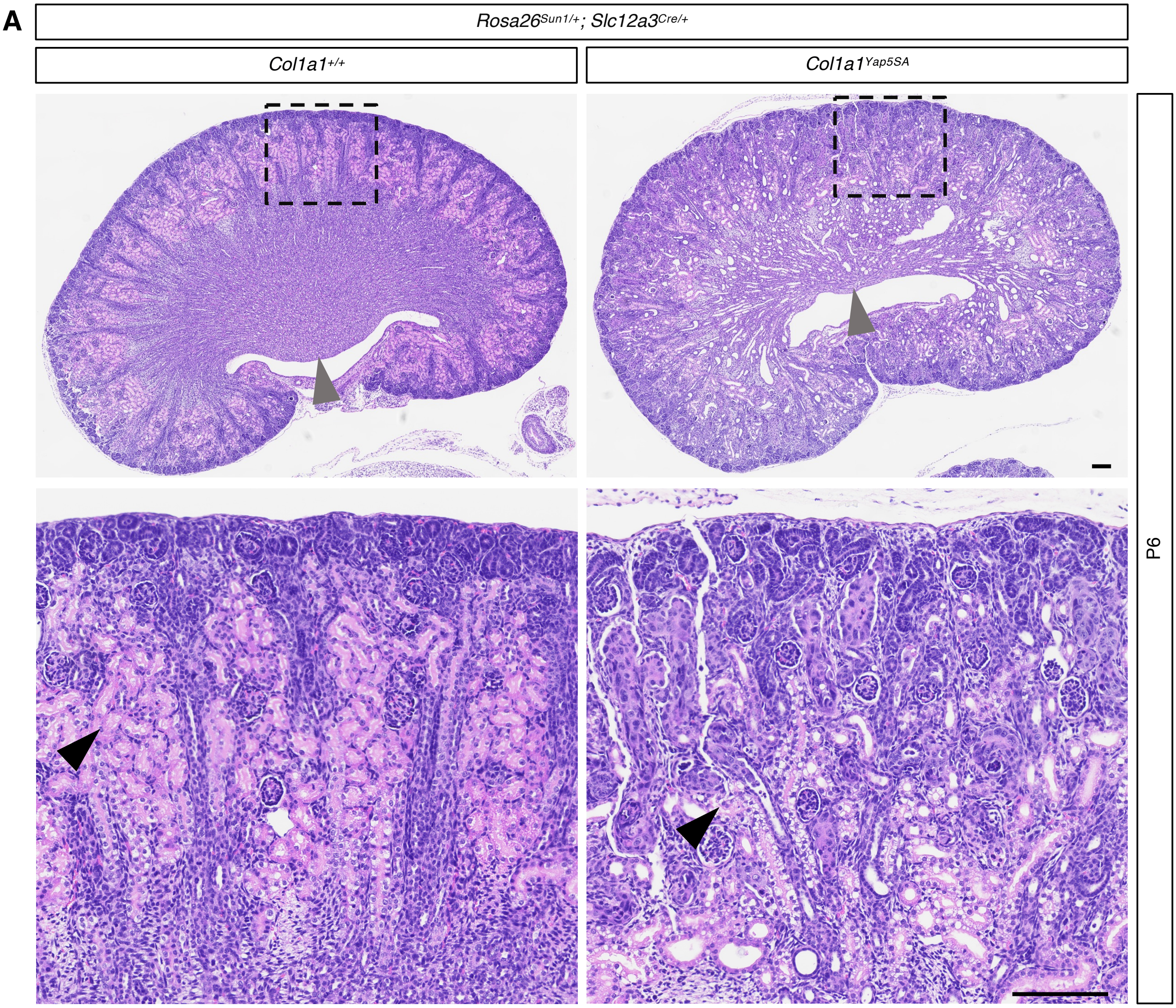


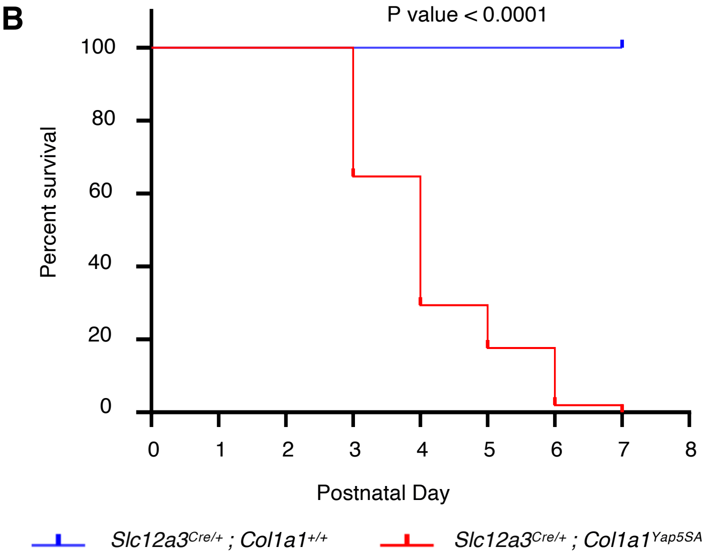


**Supplemental Figure 8. Disrupted nephron structure and postnatal lethality caused by Yap activation in the distal nephron segments**
(A) H&E staining of control and mutant kidneys. The mutant kidney shows dilation of distal nephron segments and disrupted papillary architecture (grey arrowhead). The high-magnification cortical inset (dotted squares) reveals a marked reduction in proximal tubules in the mutant (black arrowheads), indicating severe disruption of nephron architecture. Representative images from three independent experiments. Stage: P5; Scale bar: 200 µm. (B) Kaplan–Meier survival analysis showing a marked increase in neonatal mortality in mutant mice (n = 51) versus controls (n = 51).

### Supplemental Figure 9

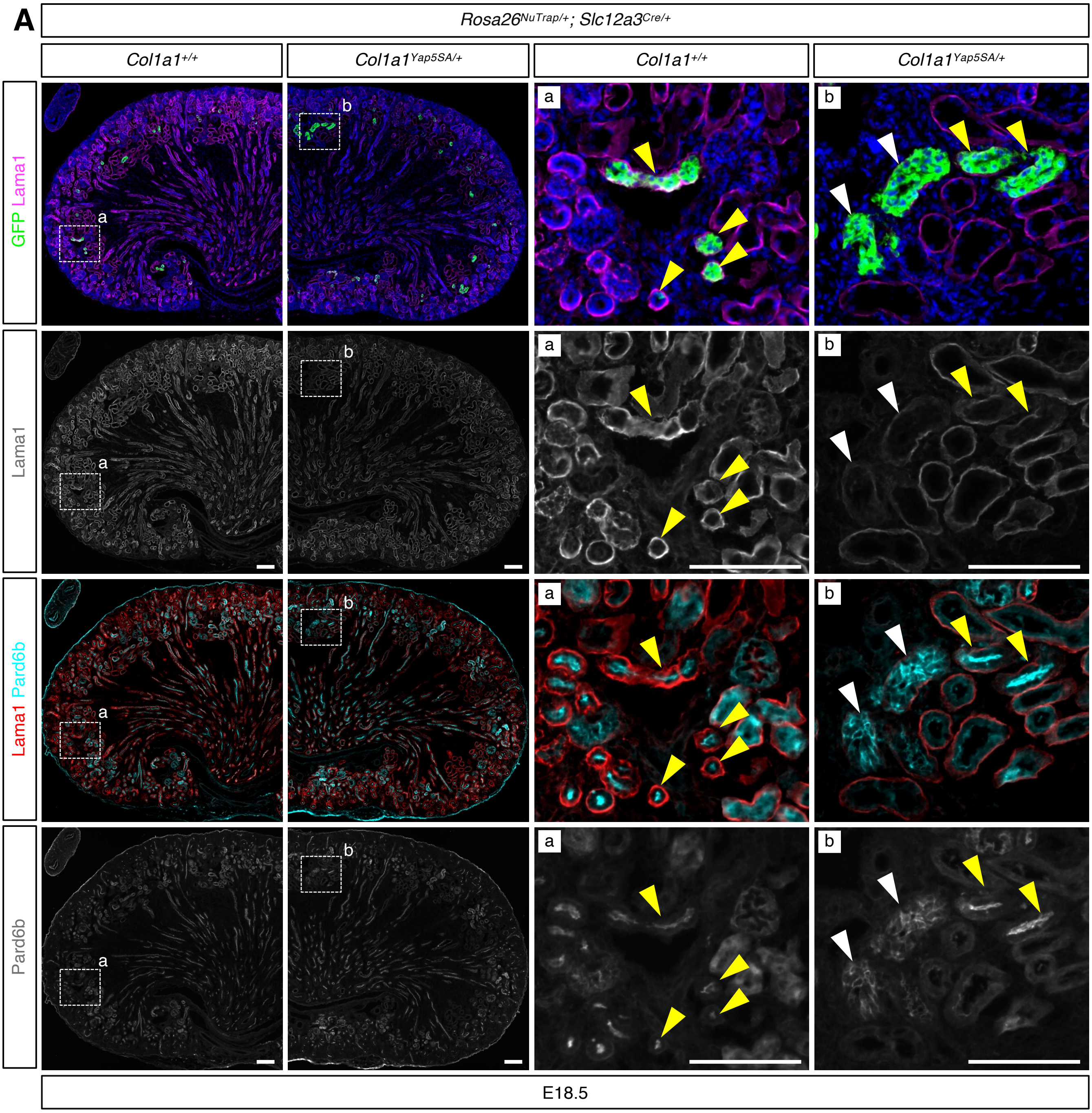

### Supplemental Figure 9

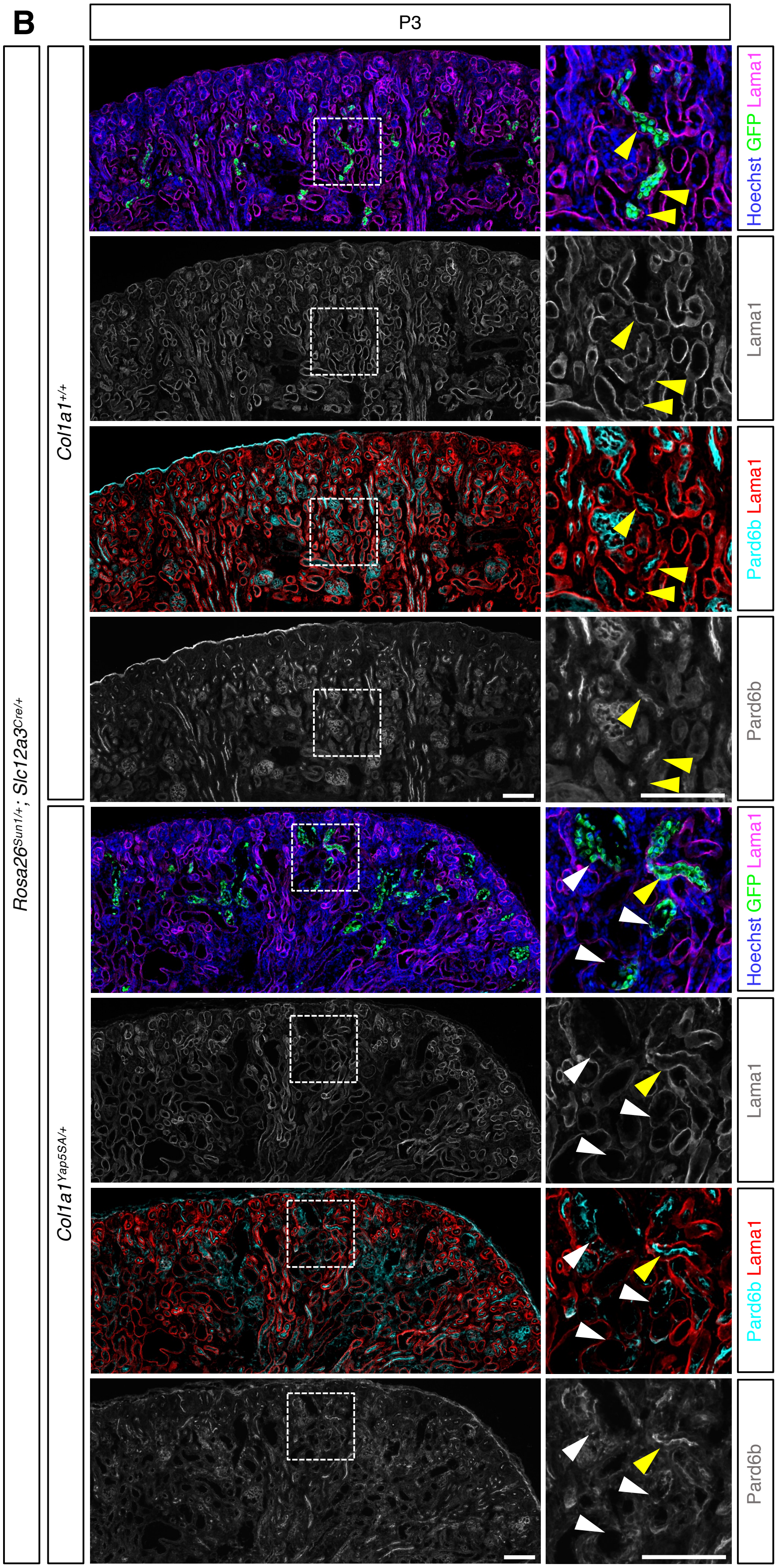


**Supplemental Figure 9. Constitutive activation of Yap in DCT and CNT leads to progressive loss of epithelial polarity.** Basal and apical polarity markers were examined at E18.5 (A) and postnatal day 3 (P3) (B). In control kidneys, GFP+ distal convoluted tubule (DCT) and connecting tubule (CNT) cells display basal Lama1 and apical Pard6b localization. In mutant kidneys, polarity defects are heterogeneous. Yellow arrowheads indicate polarity-retaining tubules, and white arrowheads indicate polarity-defective tubules. Both populations are observed at E18.5 and P3, with a higher proportion of GFP+ tubules showing loss of polarity at P3. Scale bar: 100 $\mu$m.

### Supplemental Figure 10

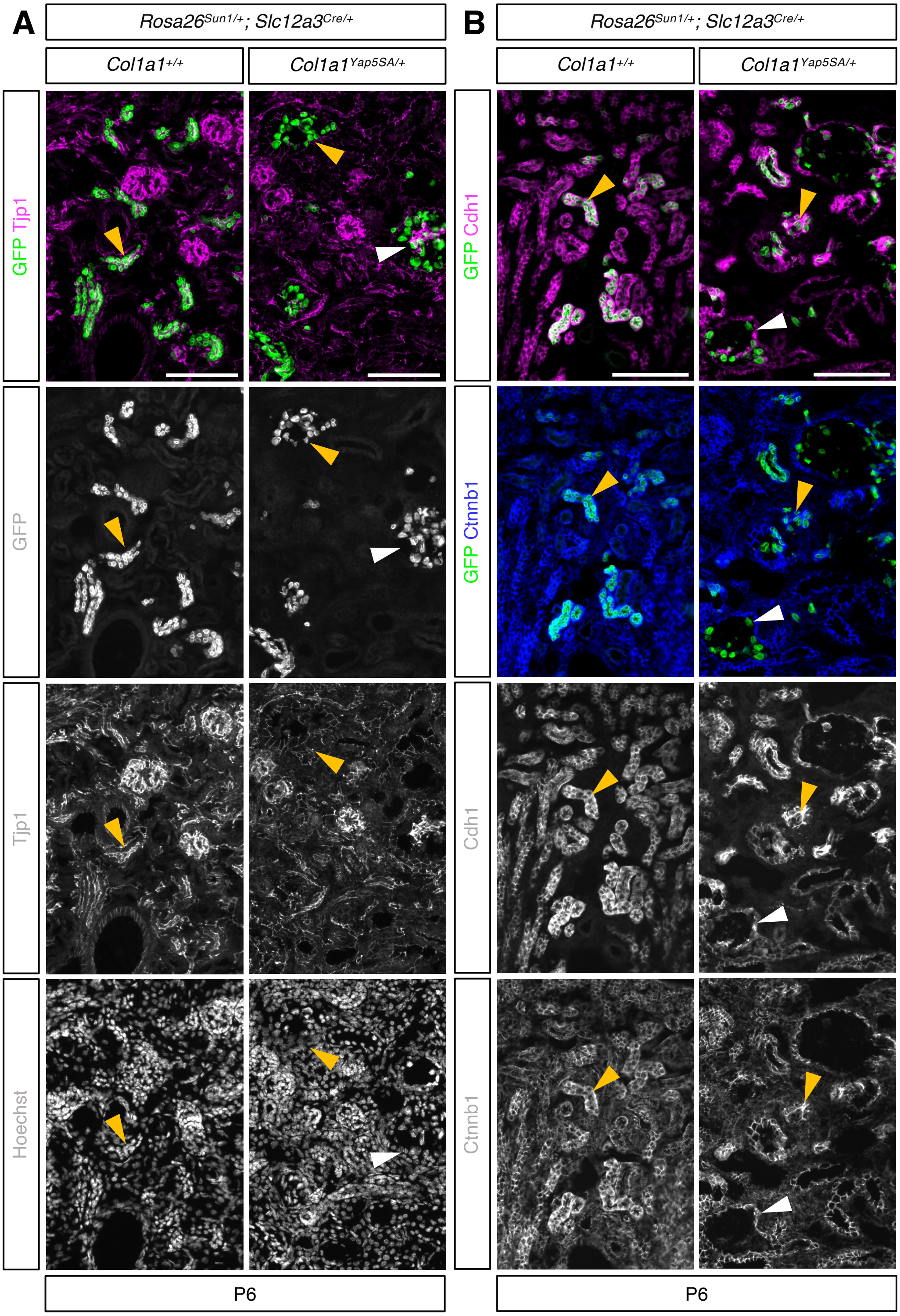


**Supplemental Figure 10. Constitutive activation of Yap in DCT and CNT impairs epithelial polarity and junctional organization** Data are identical to Figure 2C-E; colors were adjusted, and single-channel grayscale images were included to improve accessibility for readers with color-vision deficiencies. (A, B) In the control kidney, GFP+ cells express the tight junction marker *Tjp1* (*ZO1*) and the adherens junction markers, such as Ctnnb1 and Cdh1 (marked by orange arrowheads). In mutant kidneys, GFP+ cells exhibit either a loss (white arrowheads) or mislocalization (orange arrowheads) of junctional markers, suggesting impaired epithelial integrity. Representative images from three independent experiments are shown. Stage: postnatal day 6; Scale bar: 100 $\mu$m.

### Supplemental Figure 11

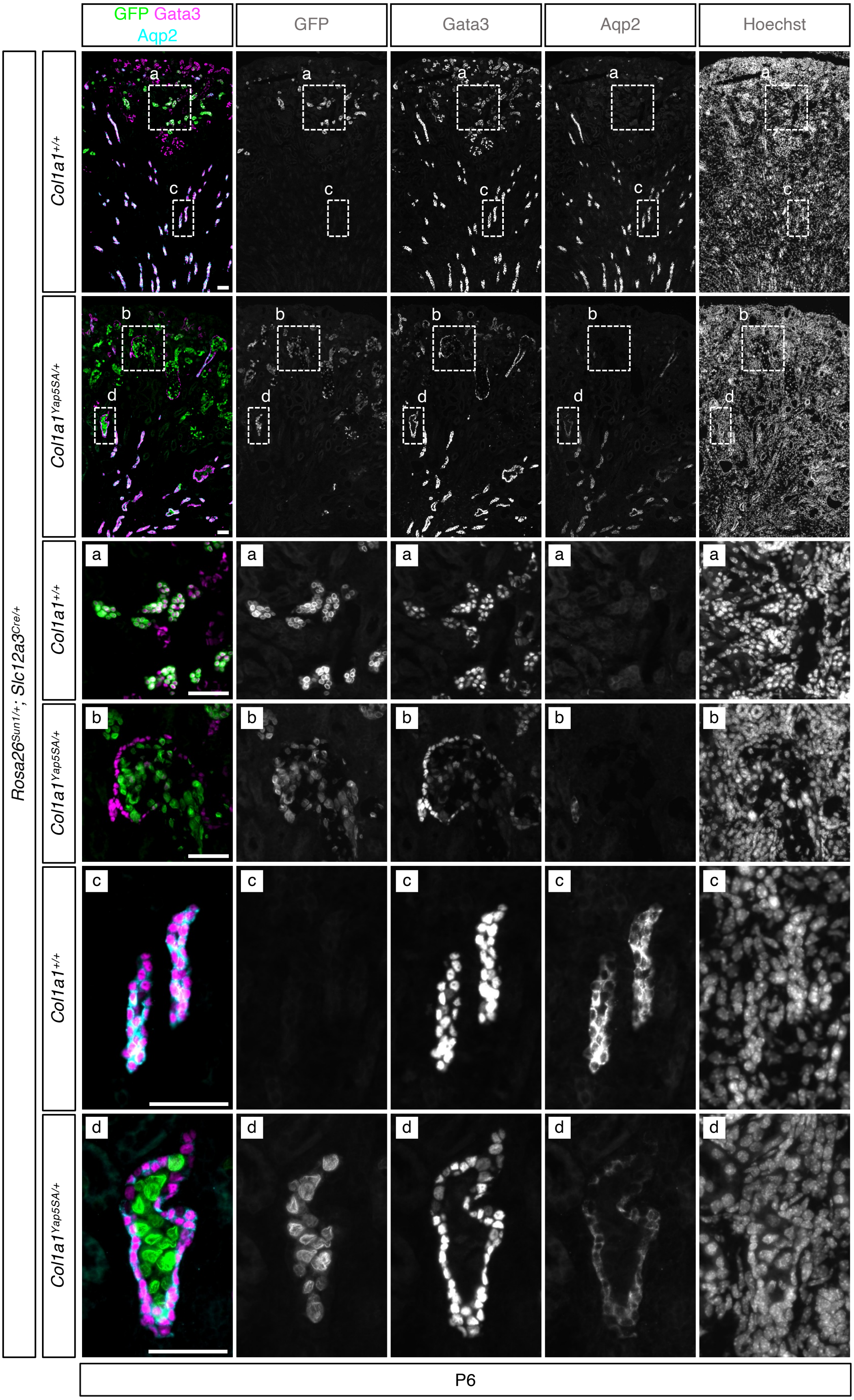

### Supplemental Figure 11

**Supplemental Figure 11. Constitutive Yap activation in DCT and CNT leads to aberrant localization of lineage-labeled cells within the collecting duct.** Data are identical to Figure 3A; colors were adjusted, and single-channel grayscale images were included to improve accessibility for readers with color-vision deficiencies. In both control and mutant kidneys, *Slc12a3Cre* activates GFP expression in DCT and CNT, but not in the collecting duct (CD). The top panels show a low-magnification view of the kidney, with boxed regions shown at higher magnification below (a–d). Aqp2 marks CD cells, whereas Gata3 marks DCT, CNT, CD, and mesangial cells. In control kidneys (a), GFP+ cells co-express Gata3. In mutant kidneys (b), GFP+ cells lack Gata3 expression. (c,d) In mutants, GFP+ cells are detected within the lumen of Aqp2+ Gata3+ CD structures. Representative images from three independent experiments are shown. Stage: postnatal day 6; Scale bar: 50 $\mu$m.

### Supplemental Figure 12

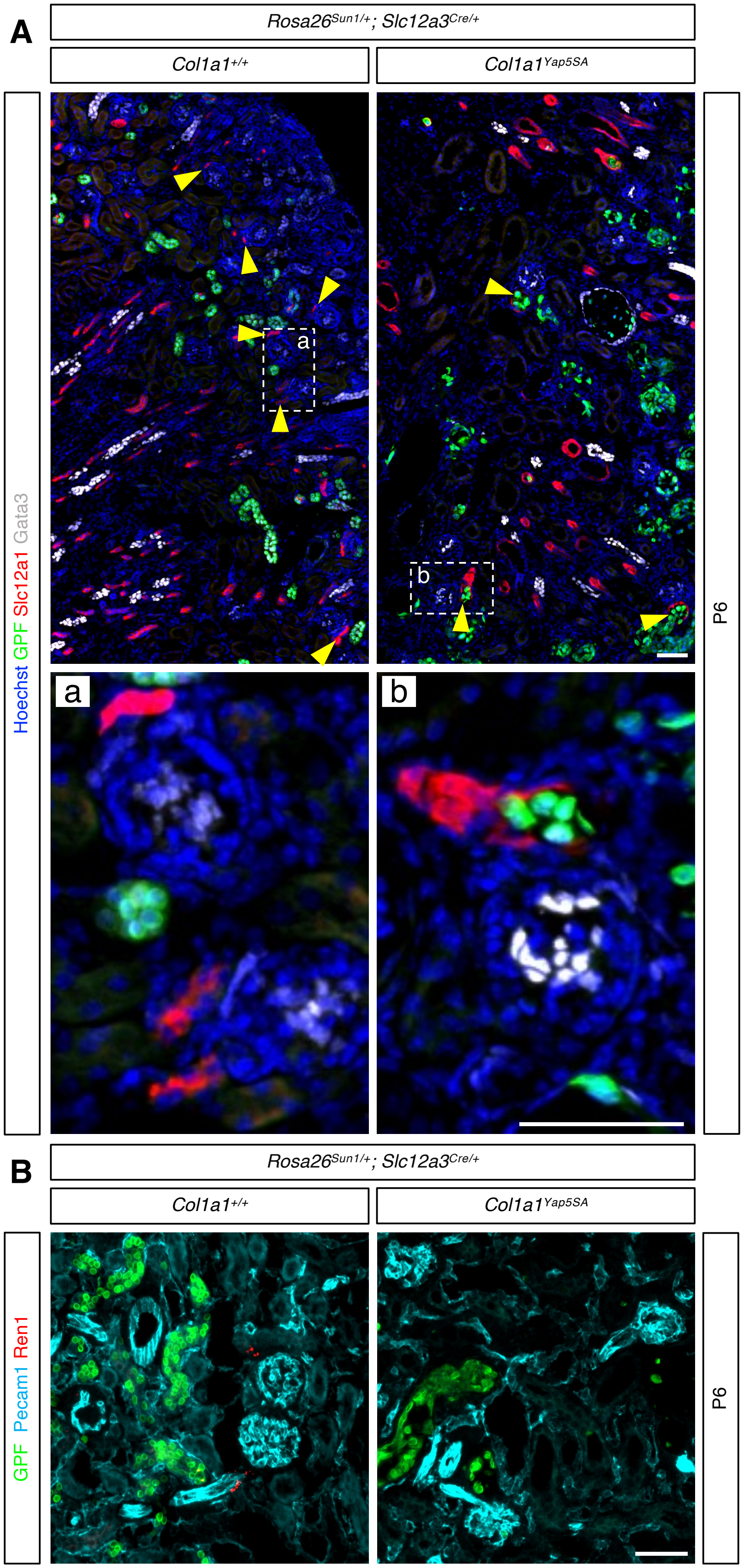


**Supplemental Figure 12. Constitutive Yap activation in distal nephron segments disrupts macula densa integrity and renin expression**

(A) Low-magnification views (upper panels) show overall kidney architecture, with the outlined regions shown at higher magnification below. Slc12a1 labels the thick ascending limb of the loop of Henle and the macula densa (MD; yellow arrowheads). Gata3 marks mesangial cells within glomeruli as well as the DCT, CNT, and collecting duct. In mutant kidneys, GFP+ Yap GOF cells are found within the lumen of Slc12a1+ MD cells (yellow arrowheads).

(B) In control kidneys, Ren1 is detected adjacent to glomeruli. In contrast, mutant kidneys exhibit complete loss of Ren1, suggesting JGA dysfunction and impaired intrarenal renin–angiotensin signaling. Pecam1 marks endothelial cells, including glomeruli. Representative images from three independent experiments. Stage: P6. Scale bar: 50 µm.

### Supplemental Figure 13

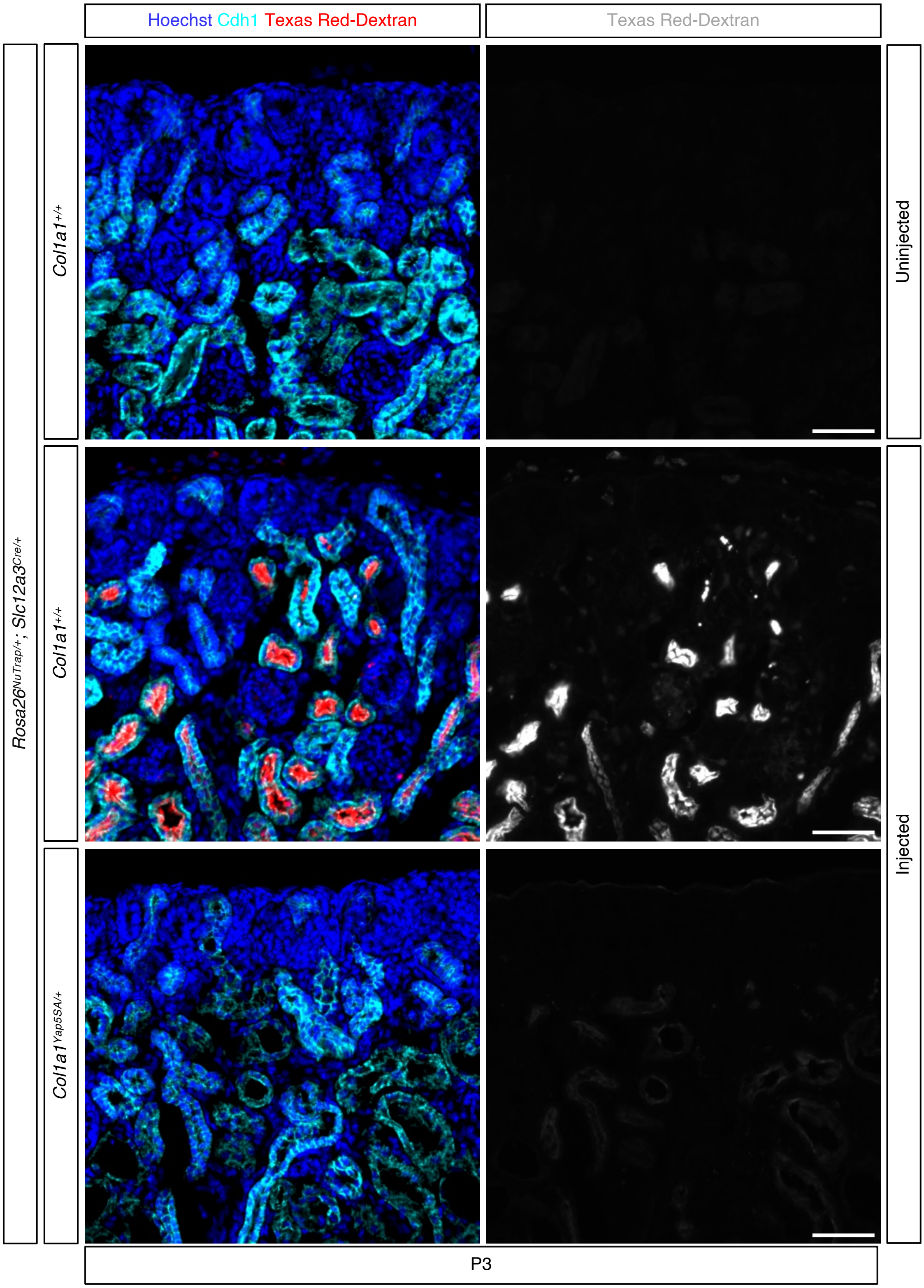


**Supplemental Figure 13. Constitutive activation of Yap in distal nephron tubules reduces luminal dextran accumulation.** In vivo dextran labeling assay in postnatal day 3 kidneys following retro-orbital injection of 10-kDa Texas Red–conjugated dextran. In control kidneys, dextran is readily detected within the lumens of Cdh1+ epithelial tubules, whereas mutant kidneys show a marked reduction in luminal dextran signal. No dextran signal is detected in uninjected controls. These results suggest that constitutive Yap activation in distal nephron segments impairs tubular luminal flow. Scale bar: 50 $\mu$m.

#### Supplemental Figure 14

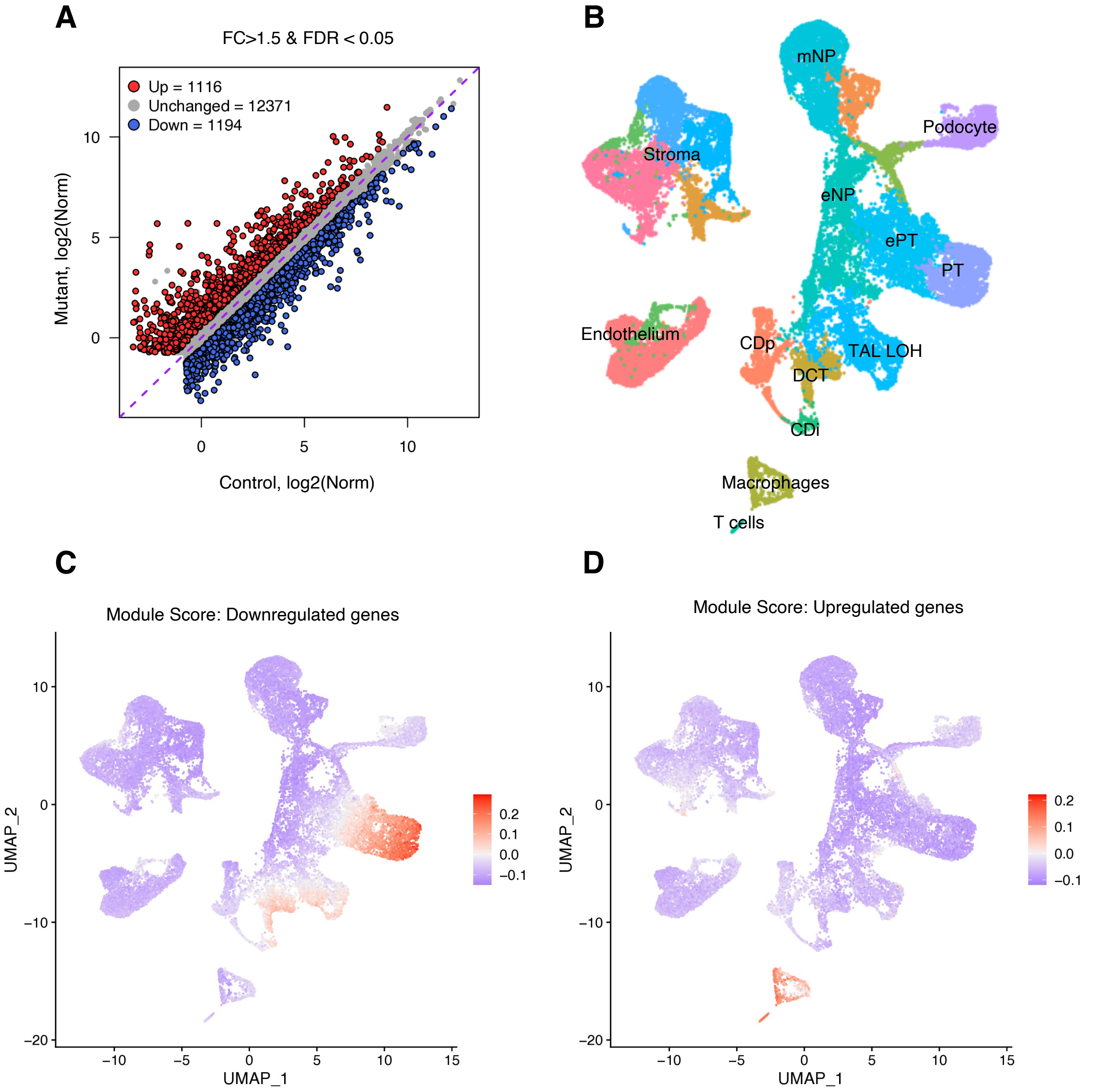


**Supplemental Figure 14. Integrating Bulk RNA-seq differential expression with single-nucleus RNA-seq to identify cell-type-specific responses to Yap activation** (A) Scatter plot displaying differentially expressed genes (DEGs) from bulk RNA-seq analysis comparing Yap GOF mutants by Slc12a3Cre to control samples. Red points indicate genes upregulated ≥2-fold with FDR < 0.05; blue points indicate downregulated genes under the same criteria; grey points represent non-significant changes. (B) UMAP plot of scRNA-seq data from mouse kidneys at E18.5 or P0 (GSE214024 and GSE275601). eNPs: epithelial nephron progenitors; mNPs: mesenchymal nephron progenitors; TAL LOH: Thick ascending limb of the loop of Henle. ePT: early proximal tubules; DCT: distal convoluted tubule; CDi: collecting duct intercalated cells; CDp: collecting duct principal cells. (C) UMAP projection showing the module score for genes downregulated in Yap GOF samples. Color intensity reflects module score: higher values indicate enrichment; lower values indicate depletion. Enrichment is concentrated in the proximal tubule cluster, suggesting Yap activation suppresses these genes in proximal tubule cells. (D) UMAP projection showing the module score for genes upregulated in Yap GOF samples. Color intensity reflects module score as defined above. Enrichment is concentrated in the clusters representing immune cells.

### Supplemental Figure 15

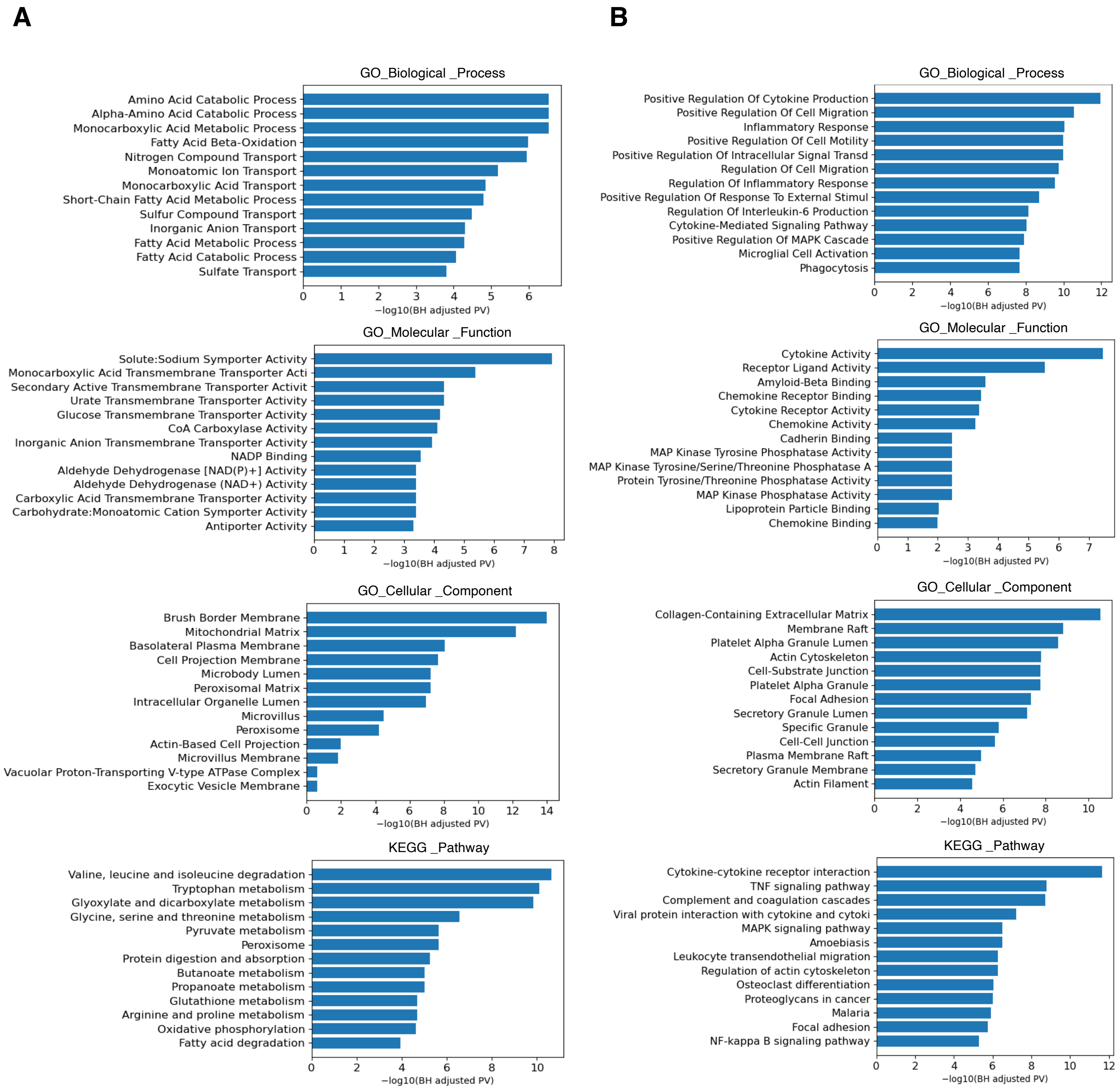


**Supplemental Figure 15. Gene Ontology and KEGG pathway enrichment analyses of differentially expressed genes in Yap GOF kidneys**

Bar plots show the top significantly enriched terms (ranked by $-\log_{10}$ adjusted P value) from Gene Ontology (GO; Biological Process, Molecular Function, and Cellular Component) and KEGG pathway analyses (adjusted $P < 0.05$). (A) Downregulated genes (n = 1,194) are enriched for metabolic and solute transport functions. (B) Upregulated genes (n = 1,116) are enriched for immune and inflammatory responses, cell migration, cytoskeletal organization, and focal adhesion signaling.

### Supplemental Figure 16

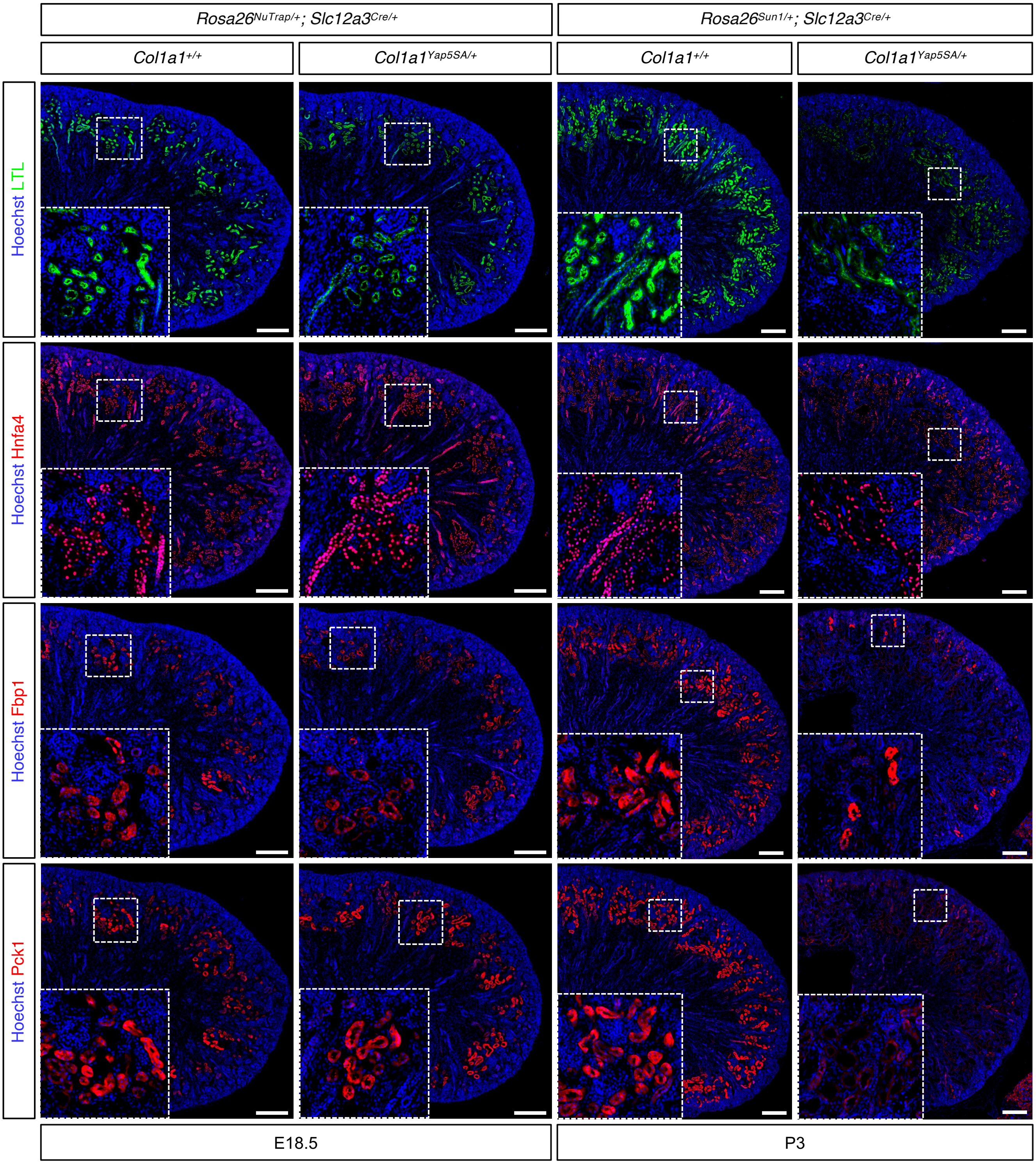


**Supplemental Figure 16. Constitutive activation of Yap in DCT and CNT affects proximal tubule development.** Proximal tubule development was assessed at E18.5 and postnatal day 3 (P3). Control kidneys show robust expression of proximal tubule markers (Hnf4a, Fbp1, and Pck1) with strong LTL (lotus tetragonolobus lectin) signal. At E18.5, mutant kidneys showed preserved expression of proximal tubule markers (Hnf4a, Fbp1, and Pck1), accompanied by weaker LTL staining. By P3, mutant kidneys show weaker LTL staining and broader reduction of proximal tubule marker expression, despite the continued presence of Hnf4a+ cells. Inset panels, indicated by white dotted squares, show higher-magnification views of the cortical regions outlined in the overview images. Scale bar: 200 $\mu$m.

### Supplemental Figure 17

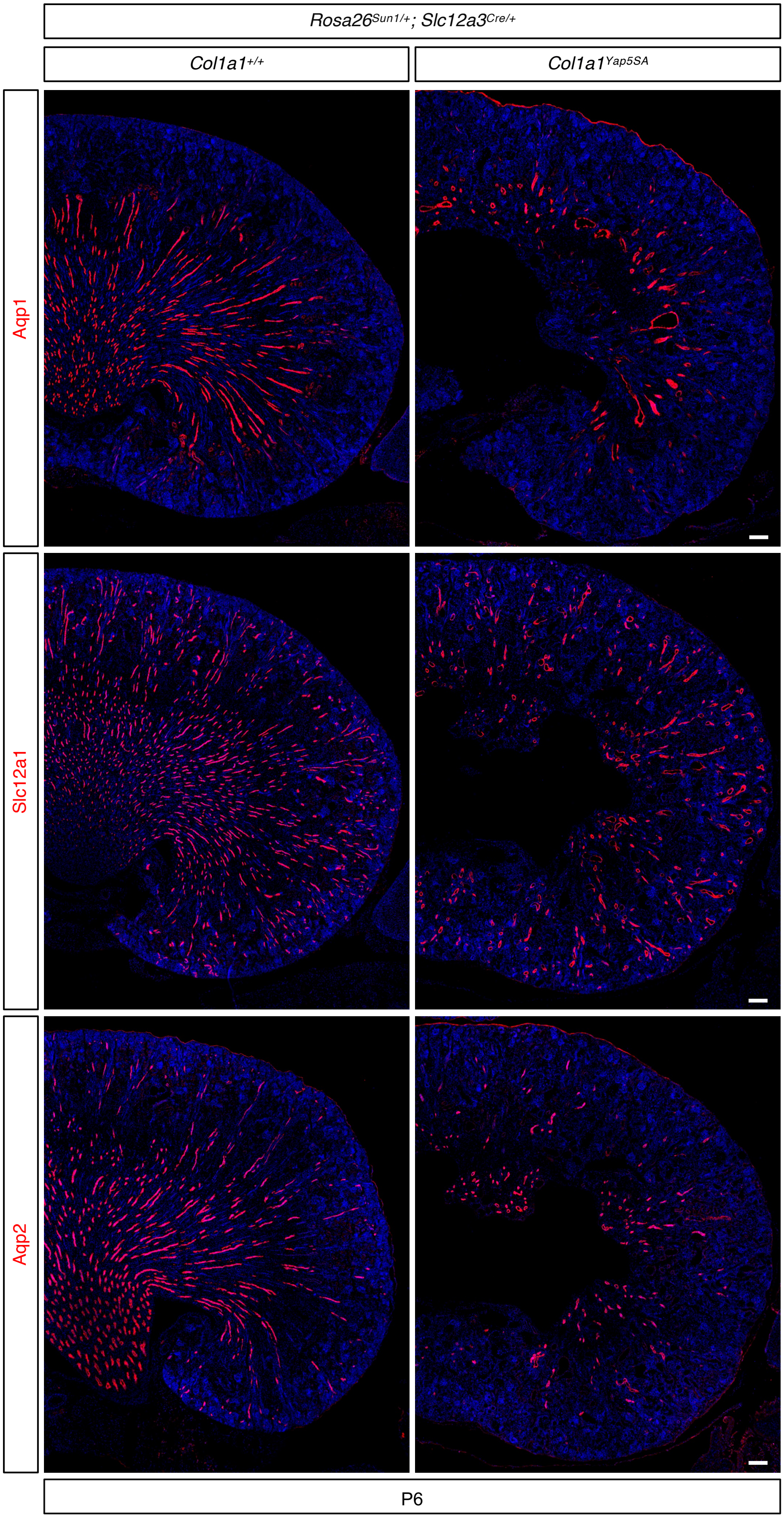


**Supplemental Figure 17. Constitutive Yap activation in distal nephron segments perturbs proper papilla formation**
Representative immunofluorescence images from control and Yap GOF kidneys showing Aqp1 (descending limb and a subset of vasa recta), Slc12a1 (thick ascending limb), and Aqp2 (collecting-duct principal cells). In control kidneys, marker expression is clearly confined to each segment, and the tubules extend in a well-organized manner. In Yap GOF kidneys, elongation of these tubular segments is markedly perturbed, and the normal spatial pattern of marker expression is disrupted, consistent with a perturbed post-natal maturation of the papilla and broad alteration of nephron patterning. Stage P6; scale bar = 100 µm.
